# Make Interactive Complex Heatmaps in R

**DOI:** 10.1101/2021.03.08.434289

**Authors:** Zuguang Gu, Daniel Hübschmann

## Abstract

Heatmap is a powerful visualization method on two-dimensional data to reveal patterns shared by subsets of rows and columns. In `R`, there are many packages that make heatmaps. Among them, **ComplexHeatmap** provides rich tools for constructing highly customizable heatmaps. It can easily establish connections between information from multiple sources by automatically concatenating and adjusting multiple heatmaps as well as complex annotations, which makes it widely applied in data analysis in various fields, especially in Bioinformatics. Nevertheless, the limit of **ComplexHeatmap** still exists. It only generates static plots which restricts deeper inspections on complex heatmaps, *e.g*., to look into a subset of rows and columns when a specific pattern of interest is observed from the heatmap. In this work, we described a new `R`/Bioconductor package **InteractiveComplexHeatmap** that brings interactivity to **ComplexHeatmap**. **InteractiveComplexHeatmap** is designed with an easy-to-use interface where static complex heatmaps can be directly exported to an interactive Shiny web application only with one extra line of code. The interactive application contains comprehensive tools for manipulating heatmaps. Besides common tools as supported in other interactive heatmap packages, **InteractiveComplexHeatmap** additionally supports, *e.g*., selecting over multiple heatmaps and searching heatmaps via row or column labels. Also, **InteractiveComplexHeatmap** provides methods for exporting static heatmaps from other popular heatmap functions, *e.g*., `heatmap.2()` or `pheatmap()`, to interactive heatmap applications. Finally, **InteractiveComplexHeatmap** provides flexible functionalities for integrating interactive heatmap widgets into other Shiny applications. **InteractiveComplexHeatmap** provides a user interface for self-defining response to the selection events on heatmaps, which helps to implement more complex Shiny web applications.

## 1. Introduction

Heatmap is a popular visualization method on two-dimensional matrices where colors are the major aesthetic elements mapping to data (Wilkinson and Friendly 2009). In routine data analysis procedures, matrix for heatmap visualization is normally accompanied with row and column reordering, *e.g*., by hierarchical clustering or seriation (Hahsler, Hornik, and Buchta 2008), so that features with similar patterns are grouped closely and they can be easily identified from the colors on heatmap. The reordering methods can also be chosen specifically for the topic under study. For example, the `R` package **EnrichedHeatmap** (Gu,Eils, Schlesner, and Ishaque 2018) can be used to visualize how DNA demethylation is enriched around the transcriptional start sites (TSS) of genes by a heatmap where columns are ordered by genomic distances from CpG dinucleotides to gene TSS and rows are ordered by the enrichment score defined by **EnrichedHeatmap**. Heatmap is widely applied in various research fields, especially popular in Bioinformatics. Since the first paper on heatmap visualization on a gene expression dataset was published (Eisen, Spellman, Brown, and Botstein 1998), heatmap has been a standard tool for visualizing *e.g*., gene expression, DNA methylation and other high-throughput datasets represented as matrices. In `R` (R Core Team 2020), there are several packages that implement heatmaps. The function `heatmap()` from package **stats** is the most fundamental one, but with very limited functionality. The function `heatmap.2()` from package **gplots** (Warnes, Bolker, Bonebakker, Gentleman, Huber, Liaw, Lumley, Maechler, Magnusson, Moeller, Schwartz, and Venables 2020) is an enhanced version of heatmap() which supports more graphics on heatmap, such as a color legend with value distribution and trace lines showing the difference of values to the column or row medians. There are also packages implemented with the **grid** graphics system such as the function `pheatmap()` from package `pheatmap` (Kolde 2019) and the function `aheatmap()` from package **NMF** (Gaujoux and Seoighe 2010). They provide more flexible controls on heatmaps.

As data emerges fast in dimensions nowadays, especially in the Genomics field, an efficient visualization for integrative analysis or multi-omics analysis is urgently needed to associate multiple types of data for easily revealing relationships between multiple objects. From the aspects of heatmap visualization, it reflects in two points. The first is the support of heatmap annotations which contain extra information to associate to the main heatmap. For example, in a typical heatmap visualization on gene expression data where rows are genes and columns are patients, it is common that patients have clinical meta-data available, such as age, gender or whether the patient has certain DNA mutations. With annotation attached to heatmap, it is easy to identify, *e.g*., whether a group of genes showing high expression correlate to a certain age interval or whether they have specific types of DNA mutations. `heatmap()` and `heatmap.2()` only support single heatmap-like annotation for one numeric or character vector. `pheatmap()` and `aheamtap()` allow multiple heatmap-like annotations for corresponding more information to heatmap. The package **superheat** (Barter and Yu 2018) and **heatmap3** (Zhao, Yin, Guo, Sheng, and Shyr 2021) support more types of graphics for annotations, such as points or lines which are able to make more accurate visual representation on annotation data. The second point of visualizing multiple sources of information is to directly apply “complex heatmap visualization” by simultaneously linking multiple heatmaps to make it straightforward to compare patterns between heatmaps. For example, in our published study (Gu et al. 2018), we applied complex heatmap visualization on gene expression, DNA methylation and various histone modifications to reveal general transcriptional regulation patterns among multiple human tissues. To implement both complex annotation and heatmap visualization, we previously developed an advanced heatmap package **ComplexHeatmap** (Gu, Eils, and Schlesner 2016a). It supports not only the basic annotation graphics as in other packages, but also a variety of extra complex annotation graphics such as violin plot or horizon plot, and it even allows users to self-define their own annotation graphics. **ComplexHeatmap** provides a simple syntax to link multiple heatmaps and annotations where rows or columns of all heatmaps are adjusted simultaneously. The simplicity of its user interface and comprehensiveness of its functionalities make **ComplexHeatmap** widely used in Bioinformatics to reveal interesting patterns from data that are potentially biologically meaningful.

After a specific pattern of subset of rows and columns is observed from the heatmap, the next step is to extract corresponding rows and columns for downstream analysis, which requires interactivity on heatmaps. There are also several `R` packages that implement interactive heatmaps such as **d3heatmap** (Cheng and Galili 2018), **heatmaply** (Galili, O’Callaghan, Sidi, and Sievert 2017) and **iheatmapr** (Schep and Kummerfeld 2017). The interactivity allows hovering on heatmap cells and selecting a region from it. There are other web-based and non-`R` implementations, such as **Morpheus** (Morpheus 2021), **Heatmapper** (Babicki, Arndt, Marcu, Liang, Grant, Maciejewski, and Wishart 2016), **Clustergrammer** (Fernandez, Gundersen, Rahman, Grimes, Rikova, Hornbeck, and Ma’ayan 2017) and **NG-CHM** (Ryan, Stucky, Wakefield, Melott, Akbani, Weinstein, and Broom 2019) which allow users to directly upload data to the web server and manipulate heatmap with no programming knowledge.

**ComplexHeatmap** is popular and powerful for generating static complex heatmaps. Here we developed a new `R`/Bioconductor package **InteractiveComplexHeatmap** that brings interactivity to **ComplexHeatmap**. **InteractiveComplexHeatmap** can be very easily applied that any static complex heatmap generated by **ComplexHeatmap** can be directly exported to an interactive Shiny web application only with one extra line of code. The interactive application contains comprehensive tools for manipulating heatmaps. Besides common tools as supported in other interactive heatmap packages, **InteractiveComplexHeatmap** additionally supports, *e.g.,* selecting over multiple heatmaps or searching heatmaps via row or column labels. The latter would be especially useful when users already have some features of interest to look into. Also, **InteractiveComplexHeatmap** provides methods to export static heatmaps from other popular heatmap functions, e.g., `heatmap.2()` or `pheatmap()`, to interactive heatmap applications, which greatly expands the ability of interactive heatmap visualization in **R**. Finally, **InteractiveComplexHeatmap** provides functionalities for integrating interactive heatmap widgets into other Shiny applications. **InteractiveComplexHeatmap** provides a user interface for self-defining response to the selection events on heatmaps, e.g., by clicking or brushing, which helps to develop more complex Shiny web applications. **InteractiveComplexHeatmap** can be freely obtained from https://bioconductor.org/packages/InteractiveComplexHeatmap/.

The paper is structured as follows. In Section 2, we first made a brief introduction to the functionalities in **ComplexHeatmap**. In Section 3, we described how to export static heatmaps to an interactive Shiny web application and we also described the tools provided by the interactive application. In Section 4, we explained how the interactivity is implemented in **InteractiveComplexHeatmap**. In Section 5, we explained in detail how to integrate **InteractiveComplexHeatmap** in Shiny application development. And finally, in Section 6, as case studies, we implemented a complex interactive heatmap application which visualizes results of differential gene expression analysis and we proposed an approach for interactively visualizing two-dimensional density distributions with heatmaps.

## 2. A brief introduction to ComplexHeatmap

In this section, we briefly introduce basic functionalities of making single heatmaps and a list of heatmaps with **ComplexHeatmap**. For more comprehensive usages, readers are recommended to refer to the **ComplexHeatmap** complete reference (Gu 2021).

**ComplexHeatmap** is implemented with the **grid** graphics engine under S4 object-oriented system. There are three major classes defined in the package: a `Heatmap` class that defines a single heatmap, a `HeatmapList` class that defines a list of heatmaps, and a `HeatmapAnnotation` class that defines a list of heatmap annotations.

### 2.1. A single heatmap

The function `Heatmap()` makes a single heatmap and returns an object in `Heatmap` class. The only mandatory argument in `Heatmap()` is a matrix, either numeric or character. It provides numerous additional arguments for customizing heatmap. Besides common functionalities also available in other heatmap functions, `Heatmap()` has following major features:

- *Flexible controls of clustering*. The hierarchical clustering accompanied with heatmap can be specified in various ways, *i.e*., 1. by a predefined distance method such as `“euclidean”` or `“pearson”`, 2. by a distance function that calculates pairwise distance between two vectors or directly from a matrix, 3. by a clustering object *e.g*. a `hclust` or a `dendrogram` object or an object that can be coerced to `dendrogram` by a proper `as.dendrogram()` function, or 4. by a clustering function that takes matrix as input and returns a dendrogram object. Besides that, dendrograms can be rendered on both edges and nodes, *e.g*., to assign different colors for dendrogram branches or to add symbols on dendrogram nodes (Figure 1A).
- *Split heatmap*. Heatmap splitting is an efficient way to highlight group-wise patterns (Figure 1D). **ComplexHeatmap** provides various ways for splitting heatmap into “slices” on both rows and columns: 1. Set a number of groups for *k*-means clustering; 2. Set a categorical variable which can be a vector or a data frame, then the heatmap is split by all combinations of levels in the categorical variable; 3. If hierarchical clustering is already applied, the splitting can be specified as a single number so that `cutree()` is internally applied to split. For the first two splitting methods, if clustering is turned on, hierarchical clustering is performed within each heatmap slice, also a second clustering is performed over heatmap slices based on the slice means.
- *Flexible controls of colors and legends*. It allows exact mapping between colors and values in the matrix with a color mapping function by specifying breaks and corresponding colors, then remaining colors are linearly interpolated in the corresponding intervals. It also allows flexible configurations on heatmap legends, such as multiple-color scheme legends. Please refer to Section “Legends” in **ComplexHeatmap** complete reference (Gu 2021) for more examples.
- *Render heatmap body as a raster image*. For a heatmap on a huge matrix that is saved as vector graphics (e.g., a `pdf` figure), rasterization helps to efficiently reduce the final file size while the loss of figure quality is ignorable. `Heatmap()` supports various methods for image rasterization. See detailed explanations and comparisons in (Gu 2020).
- *Customize heatmap*. There are two ways for customizing heatmap with user-defined code: 1. to customize heatmap body via `cell_fun` or `layer_fun` argument to add self-defined graphics to heatmap cells when heatmap is drawing (Figure 1B), and 2. to use `decorate_*()` family functions, *e.g*., `decorate_annotation()`, to add graphics to any heatmap component after the heatmap is drawn.

**Figure 1:**
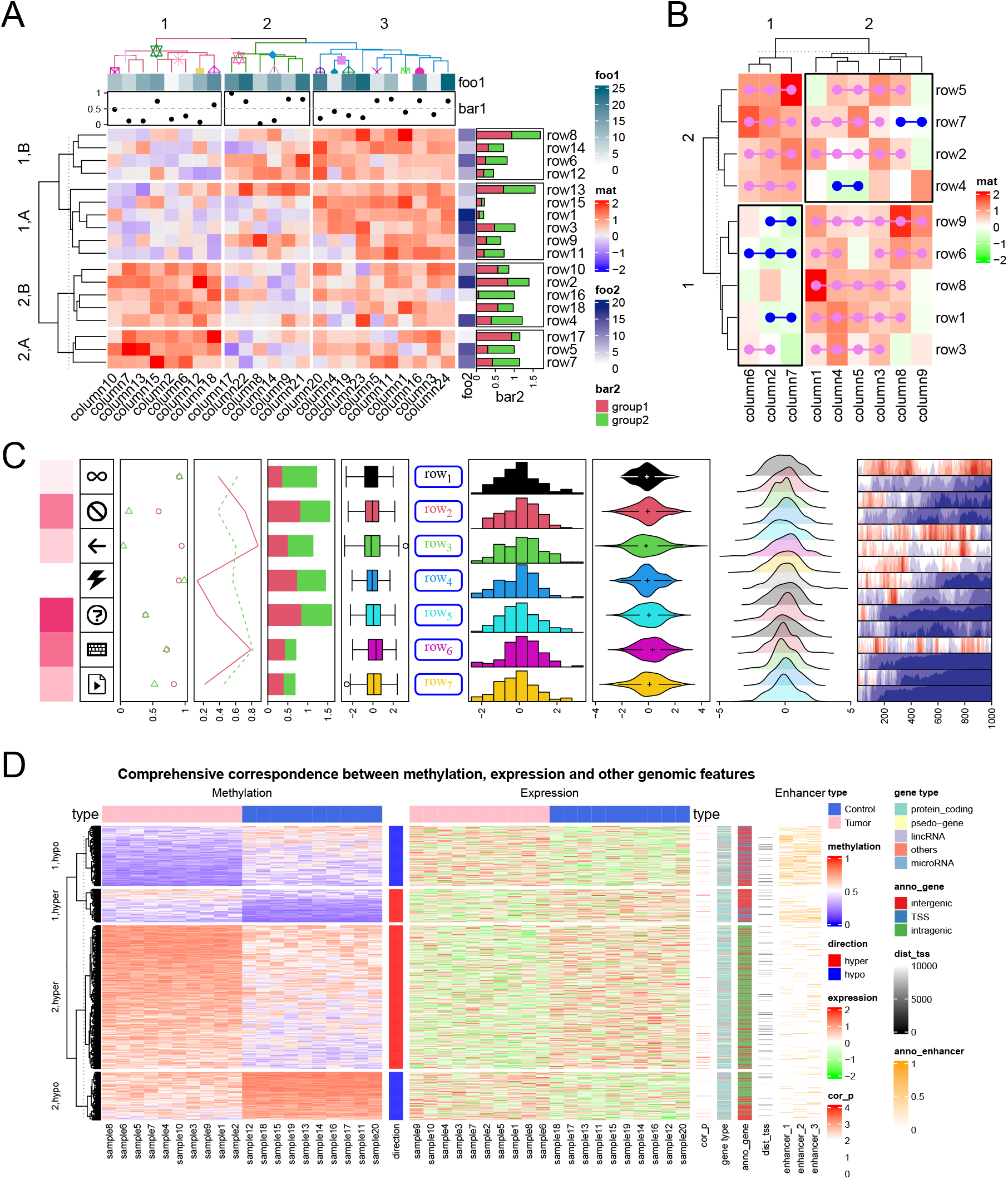
Demonstration of **ComplexHeatmap**. A) A heatmap with both row and column annotations. The columns on the heatmap were split by a three-group *k*-means clustering and rows were split by combination of a categorical variable and a two-group *k*-means clustering. B) A heatmap with customizations. On the heatmap, horizontal neighbour cells were connected if they had the same sign. Black borders were added to the top right and bottom left heatmap slices. C) Examples of various annotation graphics supported in **ComplexHeatmap**. D) An example of complex heatmap visualization based on a real-world dataset.

**Figure 2:**
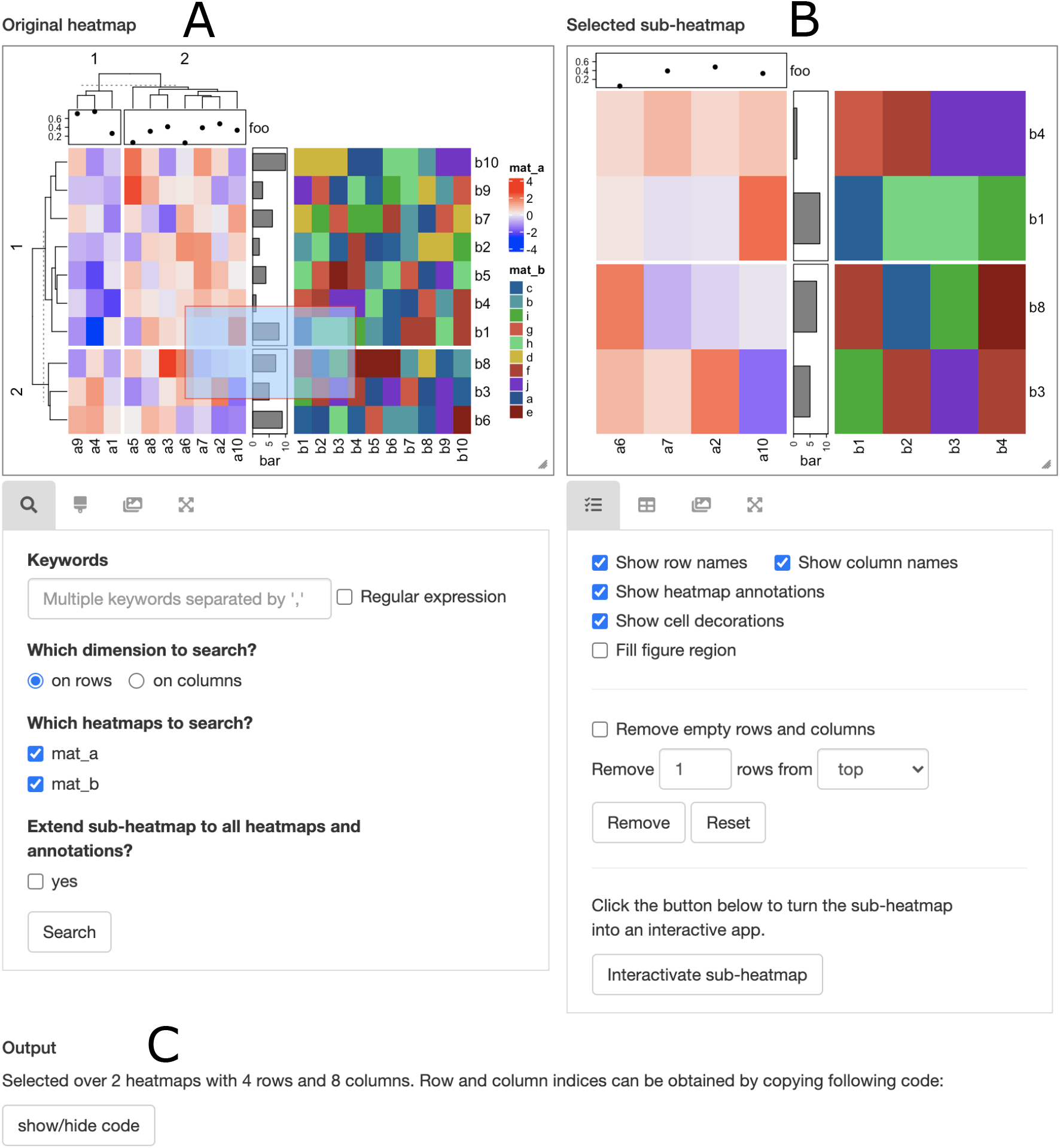
A demonstration of interactive complex heatmap. On the left is the original heatmap (labelled as “A”) and on the right is the sub-heatmap (labelled as “B”) that was selected from left. Below both heatmaps are the tools for controlling heatmaps. At the bottom of the application is the output (labelled as “C”) that shows the information of the heatmap cell that was clicked or the sub-heatmap that was selected.

### 2.2. Heatmap annotations

Heatmap annotations are important components of a heatmap that show additional information associated with rows and columns. **ComplexHeatmap** provides flexible support for annotations and defining new annotation graphics. Annotations can be put on the four sides of the heatmap and they are automatically reordered and split according to the heatmap.

There are following annotation graphics demonstrated in Figure 1C (from left to right):

- *Heatmap-like annotation*. It is named as “simple annotation” in **ComplexHeatmap**. It visualizes a vector or a matrix, either numeric or character.
- *Image annotation*. It supports images in various formats, *e.g*., `png, svg, pdf, jpg`.
- *Points annotation*. It supports a single numeric vector or a numeric matrix.
- *Lines annotation*. It supports a single numeric vector or a numeric matrix. Additionally, it supports to perform *loess* smoothing over data points.
- *Barplot annotation*. It also supports stacked bar plots.
- *Boxplot annotation*.
- *Text annotation*. It supports constructing more customized text with package **gridtext** (Wilke 2020).
- *Histogram annotation*.
- *Violin annotation*. It visualizes a list of distributions. Alternatively, the distributions can also be visualized by normal density plots or heatmaps.
- *Joy plot annotation*.
- *Horizon plot annotation*.

All built-in annotation graphics are implemented by annotation functions named with `anno_` prefix, e.g., `anno_points()` for points annotation. Besides the above listed annotations, **ComplexHeatmap** supports more complex annotations. For example, there is a “mark annotation” by `anno_mark()` which draws labels for a subset of rows or columns where the labels are shifted from their original positions to get rid of overlapping and lines are drawn to connect labels to their corresponding rows or columns. **ComplexHeatmap** provides a user interface for creating new annotation graphics. For example, package **EnrichedHeatmap** (Gu et al. 2018) defined an “enriched annotation” by `anno_enriched()` which visualizes average enrichment of certain genomic signals over a list of genomic features, and package **simplifyEnrichment** (Gu and Hübschmann 2021) defined a “word cloud annotation” by `anno_word_cloud()` which visualizes summaries of groups of rows by word clouds.

The function `HeatmapAnnotation()` accepts multiple annotations specified as name-value pairs. Simple annotations are specified as vectors, matrices or a data frame and legends are automatically generated for them. Other annotations should be specified via functions `anno_*()`. An example is as follows.

~~~
*R> ha = HeatmapAnnotation(
+     foo = runif(10),
+     bar = sample(letters[1:4], 10, replace = TRUE),
+     pt = anno_points(runif(10)),
+     txt = anno_text(month.name[1:10]),
+     …
+)*
~~~

Row annotations should be set with one additional argument `which = “row”` or by the helper function `rowAnnotation()`. Column annotations are assigned to argument `top_annotation` or `bottom_annotation` and row annotations are assigned to argument `left_annotation` or `right_annotation` in `Heatmap()`.

### 2.3. A list of heatmaps

The main feature of **ComplexHeatmap** is that it supports concatenating multiple heatmaps and annotations so that it is possible to visualize associations between various sources of information. **ComplexHeatmap** provides a simple syntax for concatenating heatmaps with the operator +. The expression returns a `HeatmapList` object and directly printing the `HeatmapList` object draws the heatmap.

~~~
*R> ht_list = Heatmap(…) + Heatmap(…) + rowAnnotation(…)
R> ht_list # It draws the heatmap*
~~~

We previously introduced annotations as components of a single heatmap. Here row annotations can also be independently concatenated to the heatmap list, as are demonstrated in the above code. Alternatively, which is less used, the heatmap lists can be vertically concatenated with the operator `%v%`.

~~~
*R> ht_list = Heatmap(…) %v% Heatmap(…) %v% HeatmapAnnotation(…)
R> ht_list*
~~~

The amount of heatmaps and annotations to be concatenated can be arbitrary. The ordering and splitting of all heatmaps are adjusted by the main heatmap, which is by default the first numeric matrix, or other heatmap in the list that is specified by the user.

The function `draw()` should be explicitly applied to the heatmap object when the heatmap is drawn in a non-interactive environment, such as inside a function, inside a `for/if-else block`, or in an `R` script running under command line. Moreover, `draw()` provides arguments for fine-tuning heatmap, such as adding self-defined legends and adjusting spaces between heatmaps.

~~~
*R> draw(ht_list, …)*
~~~

### 2.4. High-level plots implemented with ComplexHeatmap

The flexibility of **ComplexHeatmap** allows users to implement new high-level graphics on data with a matrix-like structure. In **ComplexHeatmap**, there are following high-level graphics functions that have already been implemented:

- `densityHeatmap()`. It visualizes a list of distributions via heatmap where the density values are mapped to colors.
- `oncoPrint()`. It visualizes multiple genomic alteration events (*e.g*. single-base mutations or fragment deletions) in a list of genes and in multiple patients.
- `UpSet()`. It visualizes intersections over multiple sets. This is an enhanced version of package **UpSetR** (Conway, Lex, and Gehlenborg 2017). `UpSet()` is also able to visualize intersections over genomic intervals.

More importantly, the three functions all return `Heatmap` objects, thus, the graphics can be concatenated to additional heatmaps or annotations to construct a complex view of the data. As an example, an oncoPrint can be concatenated to a gene expression heatmap to quickly connect the relationship between DNA mutations and the influence on gene expression. Example code is simply as follows:

~~~
*R> oncoPrint(…) + Heatmap(…)*
~~~

### 2.5. A complex example

Figure 1D demonstrates complex heatmap visualization on a dataset randomly generated but based on patterns found in an unpublished work. The figure visualizes associations between DNA methylation, gene expression, enhancers and gene-related information. In heatmaps, each row corresponds to a differentially methylated region (DMR, which is a genomic region showing significantly different methylation between tumor and control samples) or other objects that are associated with the corresponding DMR. In Figure 1D, there are following heatmaps from left to right:

1. A heatmap of methylation levels in DMRs. Red corresponds to 100% methylation in the DMR and blue corresponds to no methylation.
2. A one-column heatmap showing the direction of differential methylation. `hyper` means methylation is higher in tumor samples than in control and `hypo` means methylation is lower in tumor samples.
3. A heatmap of gene expression. Each gene is the nearest gene in the genome of the associated DMR. Note only the DMRs with methylation levels negatively correlated to the expression level of their nearest genes are kept in Figure 1D.
4. A one-column heatmap of p-values from Pearson correlation test between gene expression and methylation.
5. A one-column heatmap of the gene type, *i.e*., whether the corresponding gene is a protein-coding gene or of other type?
6. A one-column heatmap of the annotation of DMRs to genes, *i.e*., does the DMR locate in gene TSS or outside gene regions?
7. A one-column heatmap of the genomic distance between DMR to TSS of the associated gene.
8. A heatmap showing the overlap between enhancers and DMRs. Enhancers are genomic regions where proteins can bind to regulate gene expression.

The heatmap list is split by the combination of directions of differential methylation and a two-group *k*-means clustering. The latter is to distinguish high-methylation group and low methylation group. In Figure 1D, the complex heatmaps reveal that highly methylated DMRs are enriched in intergenic and intragenic regions and rarely overlap with enhancers (row group `“2,hypo”` and `“2,hyper”`), while in contrast, lowly methylated DMRs are enriched in TSS and enhancers (row group `“1,hypo”` and `“1,hyper”`). This might imply that enhancers associate with low methylation and methylation changes in enhancers might affect their transcriptional activities on related genes.

## 3. Export heatmap to a Shiny web application

### 3.1. The function htShiny()

For any single heatmap or a list of heatmaps represented as a `Heatmap` or a `HeatmapList` object produced by **ComplexHeatmap**, the function `htShiny()` in **InteractiveComplexHeatmap** can be simply applied to export it to a Shiny web application (Chang, Cheng, Allaire, Sievert, Schloerke, Xie, Allen, McPherson, Dipert, and Borges 2021). To demonstrate the usage, we generated a list of two heatmaps of a numeric heatmap and a character heatmap. In the numeric heatmap, *k*-means clustering with two groups was applied on both rows and columns. A point annotation was put on top of the first heatmap as a heatmap component and a barplot annotation was inserted between the first and the second heatmaps.

~~~
*R> library(ComplexHeatmap)
R> set.seed(111)
R> mat1 = matrix(rnorm(100), 10)
R> rownames(mat1) = colnames(mat1) = paste0(“a”, 1:10)
R> mat2 = matrix(sample(letters[1:10], 100, replace = TRUE), 10)
R> rownames(mat2) = colnames(mat2) = paste0(“b”, 1:10)
R>
R> ht_list = Heatmap(mat1, name = “mat_a”, row_km = 2, column_km = 2,
+        top_annotation = HeatmapAnnotation(foo = anno_points(runif(10)))) +
+     rowAnnotation(bar = anno_barplot(sample(10, 10))) +
+     Heatmap(mat2, name = “mat_b”)*
~~~

After the heatmap object `ht_list` is created, it can be directly sent to `htShiny()`. The interactive heatmap application is automatically opened in a web browser or in a pop-up window in RStudio IDE (Figure 2).

~~~
*R> ht_list = draw(ht_list) # not necessary, but recommended
R>
R> library(InteractiveComplexHeatmap)
R> htShiny(ht_list)*
~~~

To use `htShiny()`, the heatmap object is recommended to be updated with the function `draw()`. If it has not been updated, it will be applied inside `htShiny()` automatically. Updating by `draw()` speeds up loading the Shiny application because `draw()` applies clusterings which is normally the most time-consuming step in heatmap generation. After `draw()` is executed, clustering results are saved in the returned heatmap object so that repeatedly drawing heatmap can directly use the saved clustering results. If the heatmap includes randomness, such as *k*-means clustering by setting `row_km` or `column_km` argument, it is necessary to execute `draw()` before sending the heatmap object to `htShiny()` to get rid of obtaining different heatmaps when executing `htShiny()` multiple times.

Executing the function `htShinyExample()` with no argument prints a list of 54 examples of various usages of interactive heatmaps. Specifying an index in `htShinyExample()` runs the corresponding interactive application as well as showing the source code that generates the application, *e.g*., `htShinyExample(1.5)` demonstrates an example of exporting a list of two heatmaps to an interactive web application.

### 3.2 Tools in the interactive heatmap application

In the Shiny application illustrated in Figure 2, there are three main components: the original heatmap (labelled as “A” in Figure 2), the selected sub-heatmap (labelled as “B” in Figure 2) and the output (labelled as “C” in Figure 2) that prints information of the heatmap cell that was clicked or the sub-heatmap that was selected. In the original heatmap, users can click on it or select an area from it. If an area is selected from the original heatmap, a sub-heatmap is drawn on the right side of the application. Both heatmaps can be resized by dragging from their bottom right.

Under the original heatmap, there are various tools integrated:

- *Search heatmap*: If the corresponding matrices have row or column labels, it allows users to search matrix labels to obtain a subset of rows and columns. The keywords can be exactly matched to row or column labels or be a regular expression. Once the corresponding rows or columns are found in heatmaps, a sub-heatmap is drawn in the right sub-heatmap component.
- *Configure brush*: It configures visual style of the brush which selects an area from heatmap, *i.e*., border color and background color, border width and opacity of the brush.
- *Save image*: The main heatmap can be saved into a file in one of the three formats: `png, pdf` and `svg`.
- *Resize image*: The size of heatmap can be precisely controlled by manually providing a value for it.

Under the sub-heatmap, there are also tools for controlling the sub-heatmap.

- *Configure sub-heatmap*: There are three sections of controls. 1. Basic controls such as whether to show row or column labels, 2. Brushing on the original heatmap might not precisely capture rows or columns that users expected. Users can manually remove a certain number of rows or columns from four dimensions of the selected sub-heatmap. 3. Selected sub-heatmap can be further converted into a second interactive heatmap application. It will be introduced in more detail in. Section 3.4.
- *Export table*: The values in sub-heatmap can be viewed and exported as a text table. The table also includes values in corresponding annotations.
- *Save sub-heatmap*: The sub-heatmap can be saved into a file in one of the three formats: `png, pdf` and `svg`.
- *Resize sub-heatmap:* The size of sub-heatmap can be precisely controlled by manually providing a value for it.

At the bottom of the application, there is an output component. If a heatmap cell was clicked, the output component prints the meta information of that cell, such as the value, the row and column indices and labels. If there are annotations associated in heatmap, corresponding annotation values are also printed. If an area was selected from the original heatmap, the output component prints a runnable code that can be used to obtain row and column indices from the corresponding matrices. The output component can be self-defined to allow more complex output to respond to user’s actions on the heatmap. It will be introduced in more detail in Section 5.

### 3.3. Use last generated heatmaps

**ComplexHeatmap** is broadly used in many scripts and packages where they generate heatmaps but do not directly return `Heatmap/HeatmapList` objects. Nevertheless, this won’t affect the use of **InteractiveComplexHeatmap** because the last generated heatmap object is always automatically saved. Calling `htShiny()` without a heatmap object will automatically use the last one. We demonstrate this functionality with the package **cola** (Gu, Schlesner, and Hübschmann 2020).

**cola** heavily uses **ComplexHeatmap** to implement various customized heatmaps to visualize consensus clustering results as well as results from downstream analysis. As an example, the function `get_signatures()` extracts signatures that are significantly different between the predicted subgroups. `get_signatures()` makes a list of heatmaps for visualizing the patterns of signatures and returns a data frame of the signatures as well as various statistics for the statistical test, thus, we cannot directly interact with the heatmap object. When `get_signatures()` draws the heatmaps, the heatmap object is saved internally, then directly calling `htShiny()` without any argument converts the signature heatmap into an interactive application (Figure 3).

~~~
*R> library(cola) # from Bioconductor
R> data(golub_cola)
R> get_signatures(golub_cola[“ATC:skmeans”], k = 3) # this makes the heatmap
R> htShiny()*
~~~

**Figure 3:**
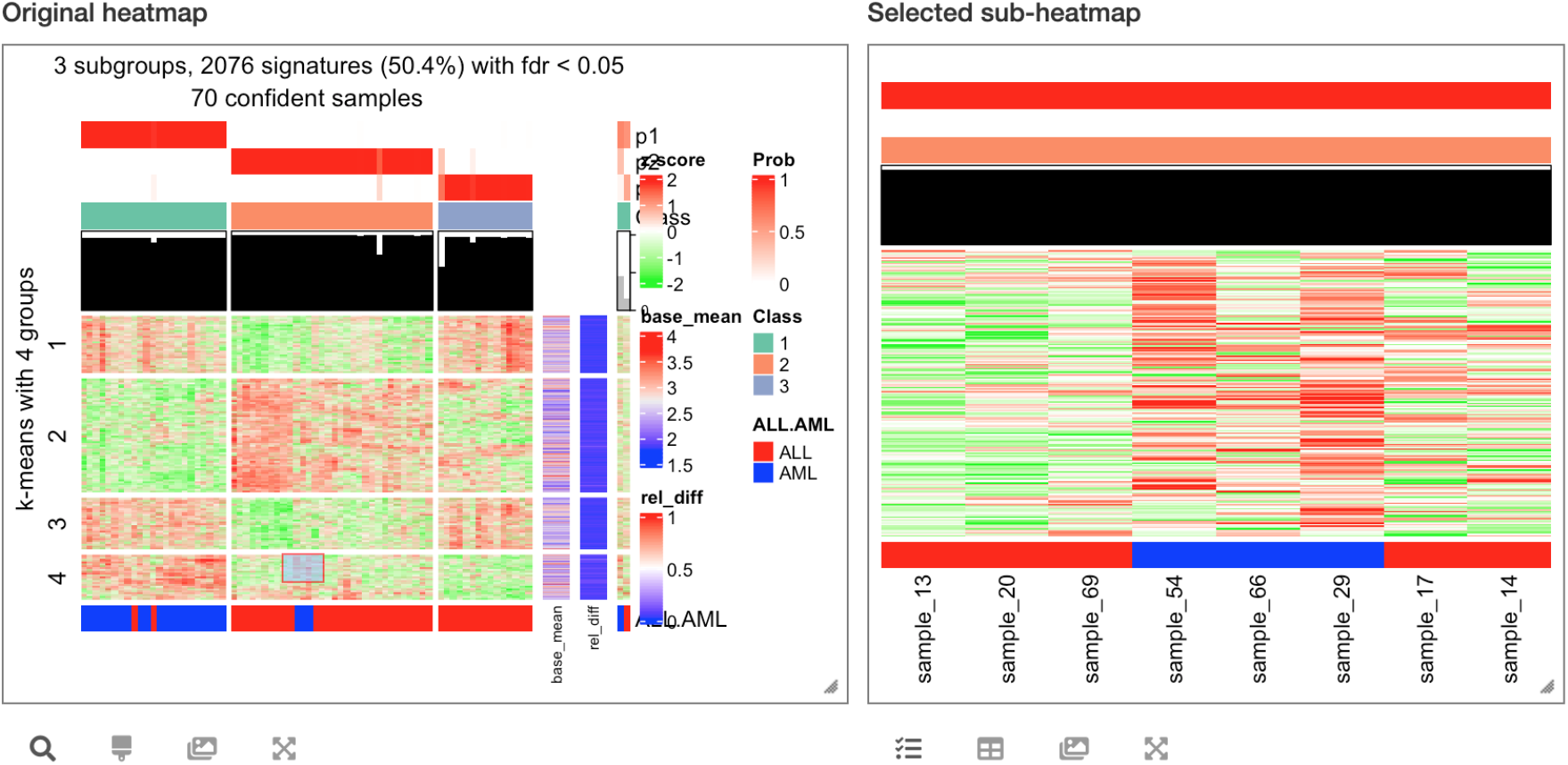
Interactivating last generated heatmap. The original heatmap is generated by function `get_signatures()` from package **cola**.

Note the functionality of automatically saving the last heatmap object is only turned on when **InteractiveComplexHeatmap** is loaded, which means, `library(InteractiveComplexHeatmap)` should be executed before making heatmap, or users can manually turn it on by executing `ComplexHeatmap::ht_opt(save_last = TRUE)`.

### 3.4. Recursive interactive heatmap application

When the original heatmap visualizes a huge matrix, in the interactive heatmap application, a small selection rectangle would cover a dense subset of rows and columns where single cells are still hard to identify in the sub-heatmap component. In this case, the sub-heatmap can be continually exported to another independent interactive heatmap widget which is in a new layer above current one, just by clicking the button “Interactivate sub-heatmap” in the tool under sub-heatmap (Figure 4). This process can be recursively applied until users are satisfied with the details seen in the sub-heatmap.

**Figure 4:**
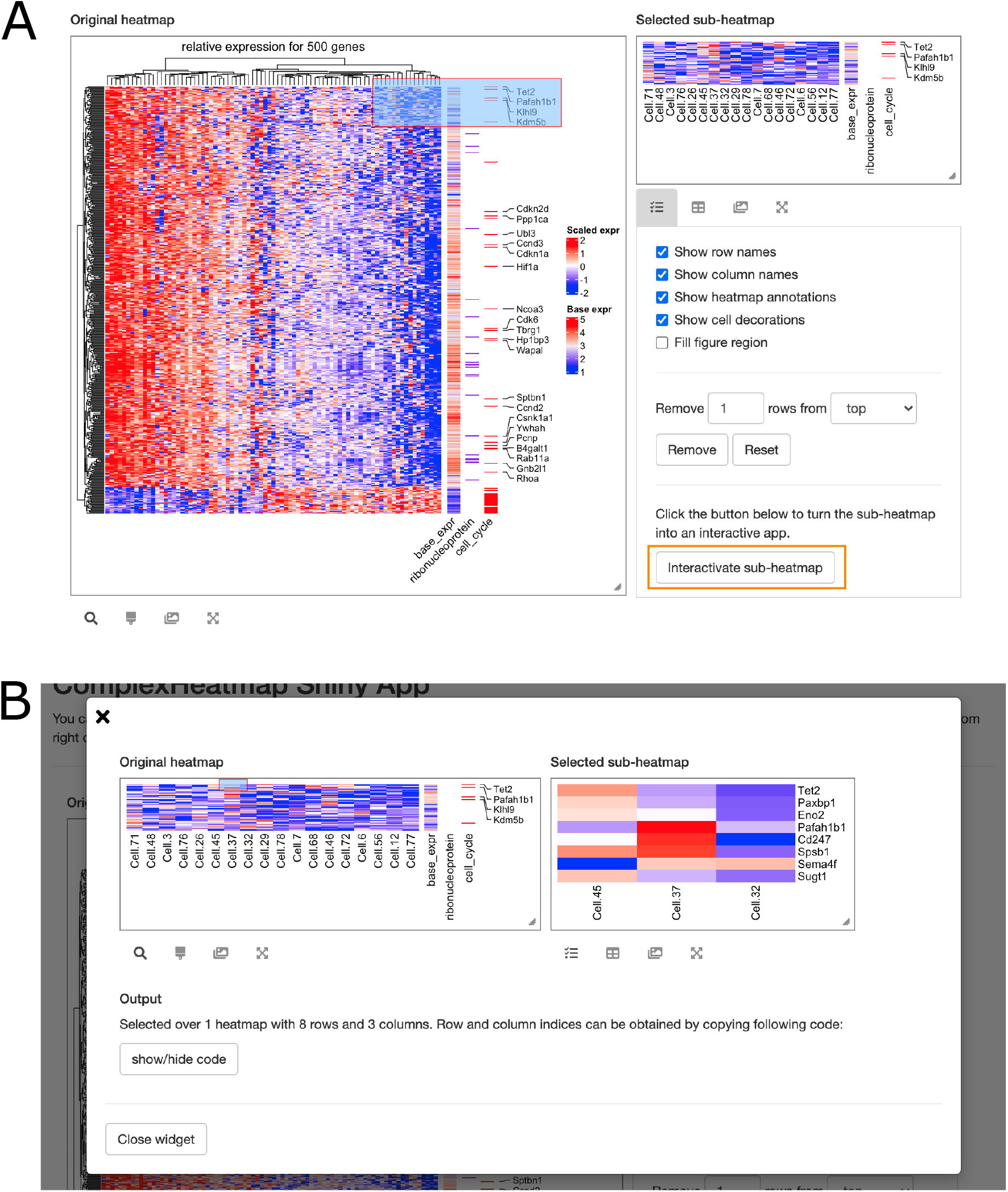
Recursively generate interactive heatmaps. The second interactive heatmap (Figure B) is generated from the sub-heatmap in the first interactive heatmap by clicking the button “Interactivate sub-heatmap” (highlighted in orange rectangle in Figure A).

### 3.5. Applications on other heatmap functions

**InteractiveComplexHeatmap** automatically interactivates static heatmaps generated by **ComplexHeatmap**. To facilitate analysis from users who still use other heatmap functions and also want to convert their heatmaps into interactive applications, in the companion package **ComplexHeatmap**, we implemented three “translation functions”: `ComplexHeatmap:::heatmap(), ComplexHeatmap:::heatmap.2()` and `ComplexHeatmap::pheatmap()` to replace `stats::heatmap()`, `gplots::heatmap.2()` and `pheatmap::pheatmap()`. The three translation functions have the same set of arguments as original ones and corresponding arguments are translated to proper settings in **ComplexHeatmap** so that they make heatmaps visually as same as possible to original heatmaps. Note in order not to dilute user’s namespace, `heatmap()` and `heatmap.2()` are not exported from **ComplexHeatmap** and users must add `ComplexHeatmap:::` prefix to use them. While for `pheatmap()`, if package **pheatmap** is already loaded ahead of **ComplexHeatmap**, there is a message printed to inform `ComplexHeatmap::pheatmap()` overwrites `pheatmap::pheatmap()`. Since the translation functions have the same sets of arguments as original ones, no further modification on code that users need to apply.

The three functions `ComplexHeatmap:::heatmap()`, `ComplexHeatmap:::heatmap.2()` and `ComplexHeatmap::pheatmap()` all generate Heatmap objects, thus heatmaps from these three functions can be converted into interactive applications. The returned object by *e.g*., `ComplexHeatmap:::heatmap()` can be sent to `htShiny()`, or `htShiny()` can be directly called with no argument after the heatmap is drawn. Example code is as follows:

~~~
*R> ht = ComplexHeatmap:::heatmap(…)
R> htShiny(ht)
R> # or even simpler
R> ComplexHeatmap:::heatmap(…)
R> htShiny()
R>
R> ComplexHeatmap:::heatmap.2(…)
R> htShiny()
R>
R> ComplexHeatmap::pheatmap(…)
R> htShiny()*
~~~

Demonstrations are in Figure 5 and live examples on the three heatmap functions can be obtained from `htShinyExample(2.4)`, `htShinyExample(2.5)` and `htShinyExample(2.6)`.

**Figure 5:**
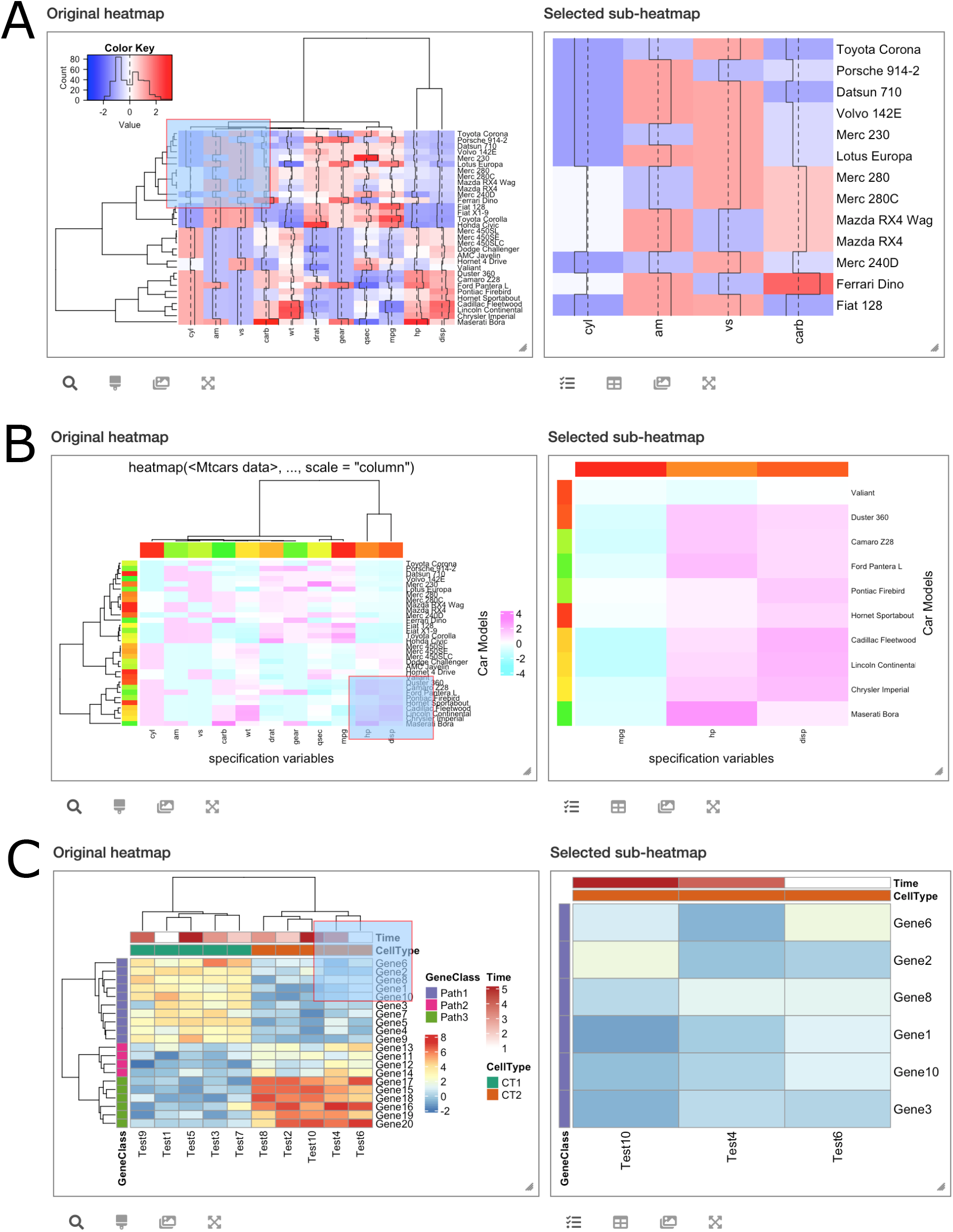
Applications on heatmaps generated from other heatmap functions. The three interactive heatmaps are from A) `heatmap()`, B) `heatmap.2()` and C) `pheatmap()`.

### 3.6. Applications on other high-level plots

**ComplexHeatmap** supports implementing high-level plots as long as the data is represented in a matrix-like structure. The following listed functions all generate Heatmap objects, thus, they can be easily exported to interactive applications.

- `ComplexHeatmap::oncoPrint()`: It visualizes various types of genomic alterations for a list of genes in multiple patients and generates a so-called *oncoPrint* (Gao, Aksoy, Dogrusoz, Dresdner, Gross, Sumer, Sun, Jacobsen, Sinha, Larsson, Cerami, Sander, and Schultz 2013). In the plot, columns are reordered in a special way to highlight the mutual exclusivity of genes that have alterations among different groups of patients. The example of an interactive oncoPrint can be found in Figure 6A and a live example can be run with `htShinyExample(2.2)`.
- `ComplexHeatmap::UpSet()`: It implements the `UpSet` plot (Conway et al. 2017) which is an efficient way for visualizing intersections in a large number of sets. Additionally, `UpSet()` is able to visualize intersections between multiple lists of genomic intervals. The example of an interactive UpSet plot can be found in Figure 6B and a live example can be run with `htShinyExample(2.3)`.
- `ComplexHeatmap::densityHeatmap()`: It visualizes a list of distributions by heatmap. The example of an interactive density heatmap can be found in Figure 6C and a live example can be run with `htShinyExample(2.1)`.
- `EnrichedHeatmap::EnrichedHeatmap()`: The package **EnrichedHeatmap** visualizes the enrichment of certain genomic signals (*e.g*., DNA methylation) on specific genomic features (*e.g*., gene TSS) by a special heatmap. **EnrichedHeatmap** inherits **ComplexHeatmap** and the constructor function `EnrichedHeatmap()` returns a Heatmap object, thus, an “enriched heatmap” can be exported to an interactive application as well. One simple example is in Figure 6D and runnable examples can be found in `htShinyExample(3.1)`, `htShinyExample(3.2)` and `htShinyExample(3.3)`.

**Figure 6:**
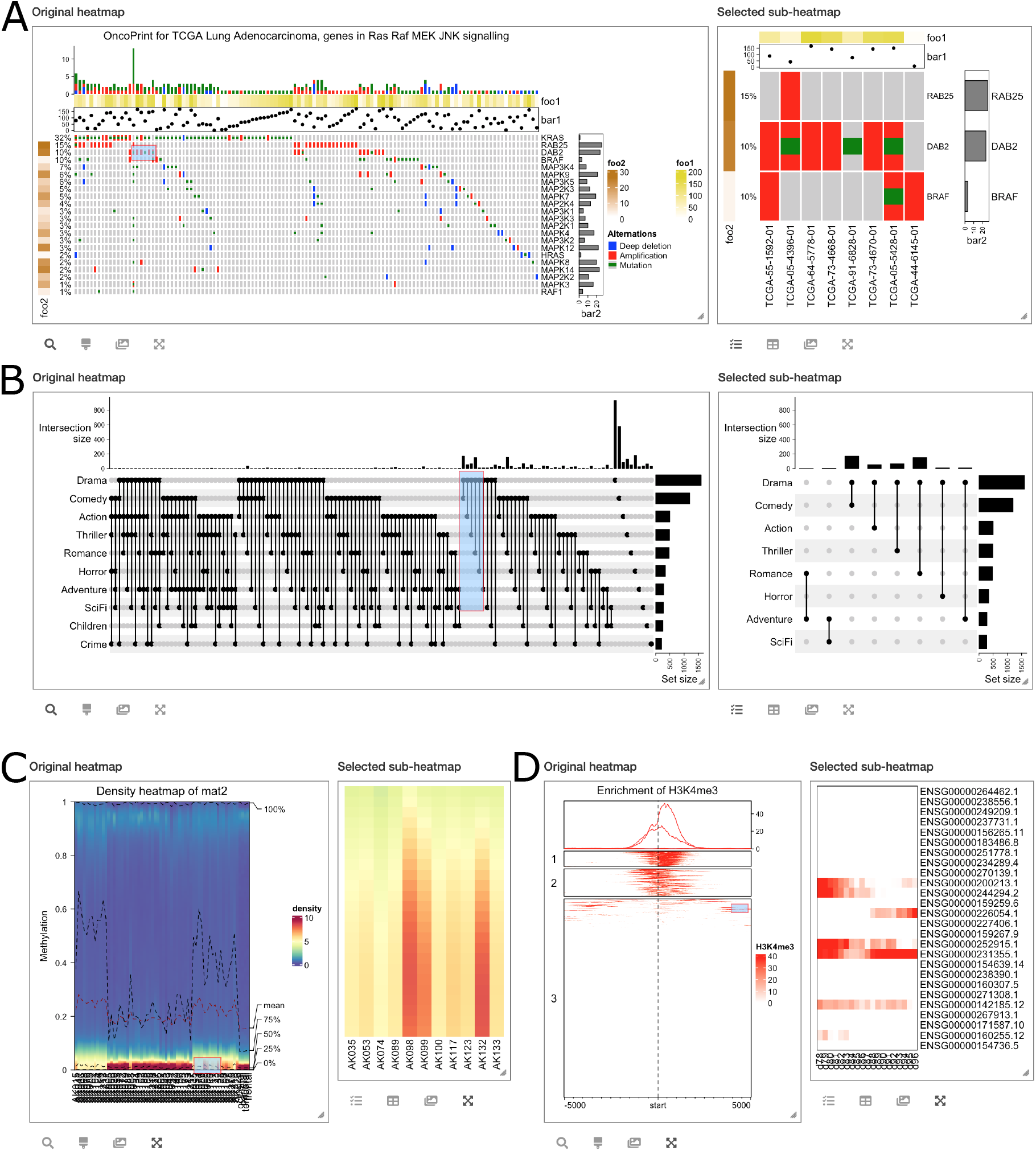
Applications on other high-level plots. A) An interactive oncoPrint. B) An interactive UpSet plot. C) An interactive density heatmap. D) An interactive enriched heatmap.

Recall **ComplexHeatmap** makes it easy to combine a list of `Heatmap` and `HeatmapAnnotation` objects, thus, heatmaps generated by the previous four functions can associate to additional heatmaps or annotations to generate complex interactive visualizations. An example is in `htShinyExample(3.2)` where the heatmap list includes an enriched heatmap of histone modification on gene TSS, a second enriched heatmap of DNA methylation on gene TSS and a third normal heatmap of gene expression; the heatmap list clearly illustrates the correspondence between three genomic signals and the interactivity of this Shiny application ensures the genes with specific patterns can be easily captured.

As we have demonstrated in Section 3.3, for some other high-level plots generated by functions which internally use **ComplexHeatmap** while do not directly return `Heatmap` or `HeatmapList` objects (*e.g*., `cola::get_signatures()`), users can always first generate heatmap in a certain graphics device (on-screen or off-screen), then execute `htShiny()` with no heatmap object to obtain the interactive applications (see examples in `htShinyExample(1.6)` and `htShinyExample(1.7)`).

### 3.7. Interactivate heatmaps indirectly generated by heatmap(), heatmap.2() and pheatmap()

In Section 3.3, we introduced how to export heatmaps indirectly generated from **ComplexHeatmap** to interactive applications. There is another scenario where heatmaps are indirectly generated by `stats::heatmap()`, `gplots::heatmap.2()` or `pheatmap::pheatmap()`, *i.e*., they are generated by third-party functions which internally use these three heatmap functions. How can we turn these heatmaps into interactive applications? The solution is simple. We just need to go to *e.g*. **pheatmap** namespace and replace `pheatmap` with `ComplexHeatmap::pheatmap` right there.

The following example is from package **SC3** (Kiselev, Kirschner, Schaub, Andrews, Yiu, Chandra, Natarajan, Reik, Barahona, Green, and Hemberg 2017) where function `sc3_plot_expression()` draws a heatmap internally using `pheatmap::pheatmap()` (This can be found out by checking the source code of `sc3_plot_expression()`).

~~~
*R> # The following code is runnable
R> library(SingleCellExperiment)
R> library(SC3)
R> library(scater)
R>
R> sce = SingleCellExperiment(
+     assays = list(counts = as.matrix(yan),
+            logcounts = log2(as.matrix(yan) + 1)),
+      colData = ann
+)
R>
R> rowData(sce)$feature_symbol = rownames(sce)
R> sce = sce[!duplicated(rowData(sce)$feature_symbol),]
R> sce = runPCA(sce)
R> sce = sc3(sce, ks = 2:4, biology = TRUE)
R>
R> sc3_plot_expression(sce, k = 3)*
~~~

To replace the internal use of `pheatmap::pheatmap()` with `ComplexHeatmap::pheatmap()`, the function `assignInNamespace()` can be used to directly change the value of `pheatmap` in **pheatmap** namespace. After doing that, executing `sc3_plot_expression()` always uses `ComplexHeatmap::pheatmap()` and now it is possible to use `htShiny()` to export it to an interactive application.

~~~
*R> assignīnNamespace(“pheatmap”, ComplexHeatmap::pheatmap, ns = “pheatmap”)
R> library(īnteractiveComplexHeatmap)
R> sc3_plot_expression(sce, k = 3)
R> htShiny()*
~~~

To safely change `stats::heatmap()`, `gplots::heatmap.2()` or `pheatmap::pheatmap()` to `ComplexHeatmap:::heatmap()`, `ComplexHeatmap:::heatmap.2()` or `ComplexHeatmap::pheatmap()`, it is recommended to execute one of the following commands at the beginning of user’s `R` session.

~~~
*R> assignInNamespace(“heatmap”, ComplexHeatmap:::heatmap, ns = “stats”)
R> assignInNamespace(“heatmap.2”, ComplexHeatmap:::heatmap.2, ns = “gplots”)
R> assignInNamespace(“pheatmap”, ComplexHeatmap::pheatmap, ns = “pheatmap”)*
~~~

## 4. Implementation of interactivity

Being different from interactive heatmap packages based on `JavaScript`, *e.g*., **d3heatmap**, **heatmaply** or **iheatmapr**, **InteractiveComplexHeatmap** has a special way to capture positions that users clicked on heatmaps and to extract values from corresponding matrices. To demonstrate it, we still use the object `ht_list` previously generated in Section 3.1 which includes a list of two heatmaps and *k*-means clustering was applied on the numeric heatmap.

**InteractiveComplexHeatmap** implements two types of interactivity: 1. on the interactive graphics device, and 2. in the Shiny web application. The interactivity on the interactive graphics device is the basis of the interactivity in the Shiny application, thus, in following sections, we will first introduce how the interactivity is implemented with the interactive graphics device.

### 4.1. On the interactive graphics device

Here the “interactive graphics device” refers to the window for generating plots if `R` is directly used in the terminal, or the figure panel in Rstudio IDE.

**InteractiveComplexHeatmap** first captures physical positions of all heatmap slices, *i.e*., the distance to the bottom left of the graphics device, by the function `htPositionsOnDevice()`. Internally, the function goes to the viewport of every heatmap slice and utilizes the function `grid::deviceLoc()` to capture positions measured in the graphics device. Before executing `htPositionsOnDevice()`, the heatmap should be already drawn on the device so that `htPositionsOnDevice()` can access various viewports.

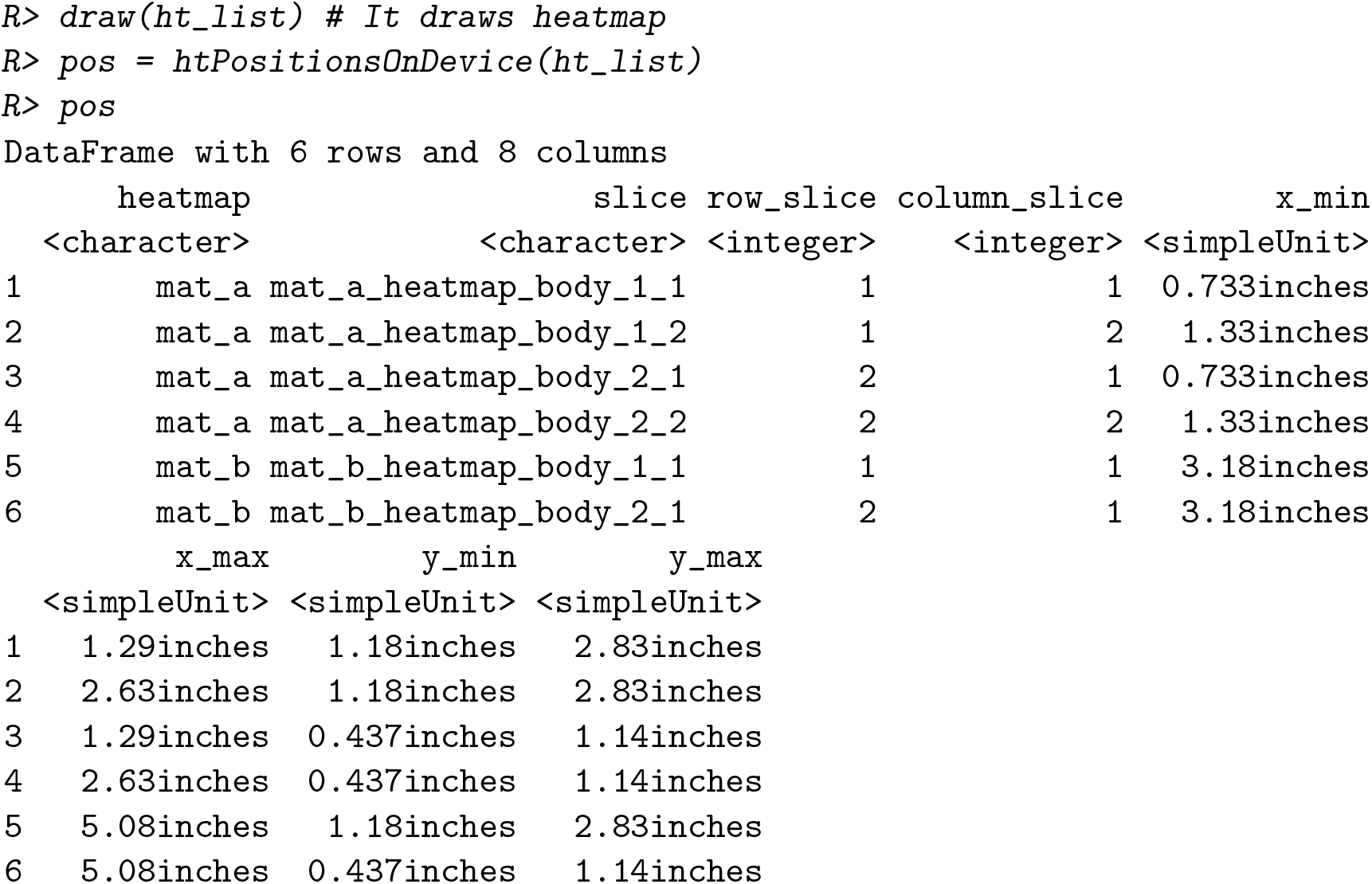

Note, the calculation of positions of heatmap slices relies on the device size. In the previous example code, pos was calculated in a device with 6 inches width and 4 inches height.

The returned object `pos` is a `DataFrame` object that contains positions of all heatmap slices. The `DataFrame` class is defined in package **S4Vectors** (Pagès, Lawrence, and Aboyoun 2020) and is very similar to a data frame, but it can store more complex data types, such as the `simpleUnit` vectors generated by `grid::unit()`. Figure 7A contains the original heatmap and black rectangles in Figure 7B were drawn based on the positions that were captured.

**Figure 7:**
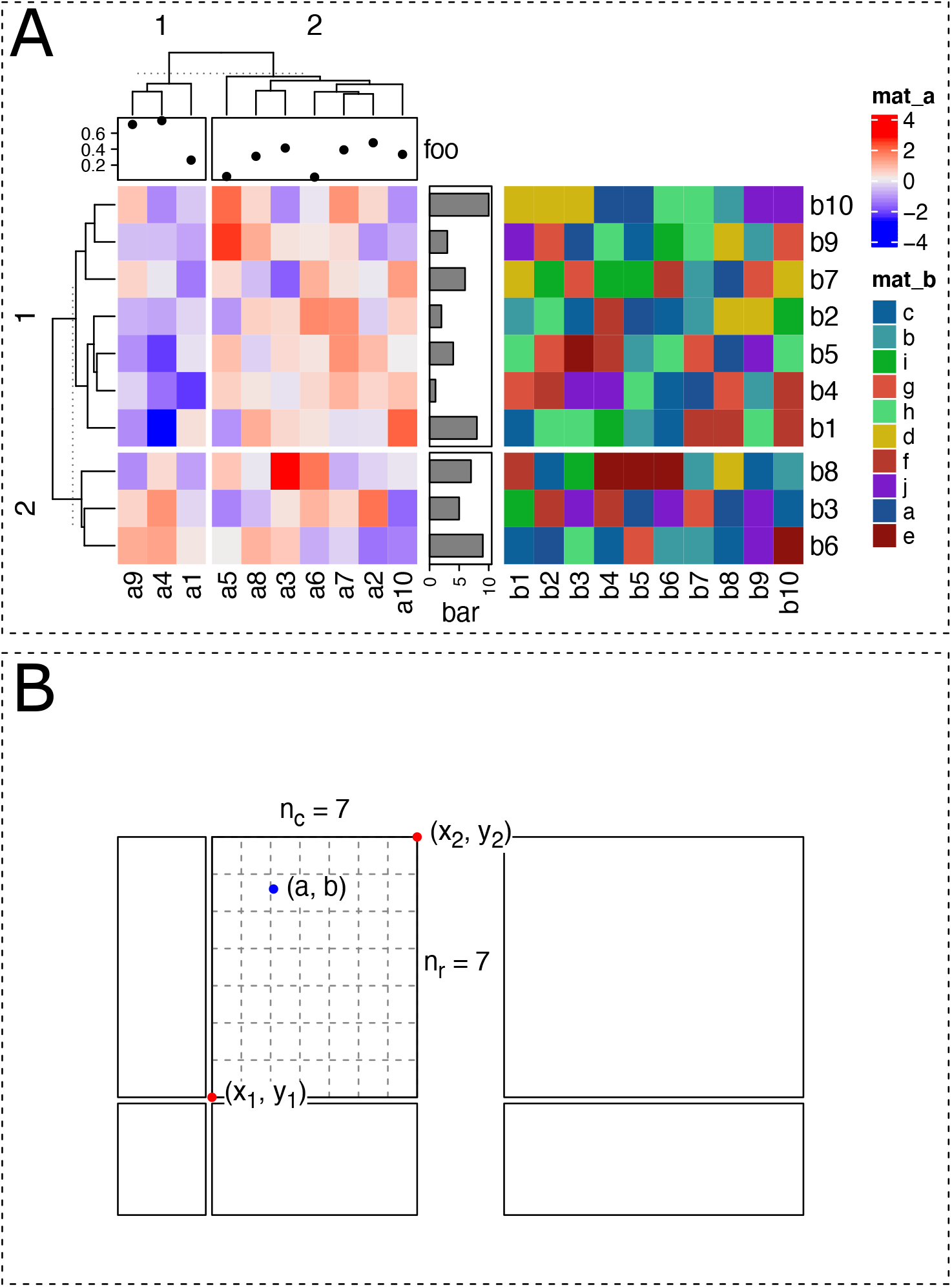
Demonstration of the implementation of interactivity. A) The original heatmap. B) The black rectangles are drawn based on the positions captured for all heatmap slices. Figure A and B were both drawn in a device with 6 inches width and 4 inches height. Dashed rectangles correspond to image borders.

Next, **InteractiveComplexHeatmap** captures the physical position of the point that the user clicked on the heatmap. This is simply done by applying function `grid::grid.locator()` in the `ROOT` viewport. Now with knowing positions of heatmap slices and position of the point that the user clicked, it is possible to calculate which row and column in the original matrix that user’s click corresponds to.

In Figure 7B, assume blue point with coordinate (*a, b*) was clicked by the user. The heatmap slice which the user clicked into can be easily identified by comparing the position of the click and the position of every heatmap slice. Assume the heatmap slice where user’s click is in has a range of (*x*_1_,*x*_2_) on *x* direction and a range of (*y*_1_,*y*_2_) on *y* direction. There are *n_r_* rows (*n_r_* = 7) and *n_c_* columns (*n_c_* = 7) in this heatmap slice and they are marked by dashed lines in Figure 7B. Note all coordinate values (*a, b*, *x*_1_, *y*_1_, *x*_2_ and *y*_2_) are measured as the distances to the bottom left of the graphics device.

In this heatmap slice, the row index *i_r_* and column index *i_c_* of the cell where the click is in can be calculated as (assume the left bottom corresponds to the index of 1 for both rows and columns):

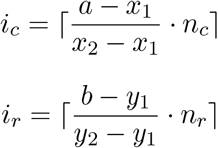

where the symbol ⌈*x*⌉ means the ceiling of the numeric value *x*. In **ComplexHeatmap**, the row with index 1 is always put on the top of the heatmap, then *i_r_* should be adjusted to:

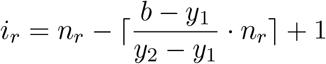

The subset of row and column indices in the original matrix that belongs to the selected heatmap slice is already stored in `ht_list` object (they can be retrieved by functions `row_order()` and `column_order()`), thus, we can obtain the row and column index in the original matrix that corresponds to user’s click easily with i_r_ and *i_c_*.

Denote the matrix for the complete heatmap (without splitting) as *M*, and denote the subset of row and column indices in the heatmap slice where user’s click is in as *o*^row^ and *o*^col^. Note, *o*^row^ and *o*^col^ can be reordered due to *e.g*. clustering. Then row and column indices for the clicked point in *M*, denoted as *j_r_* and *j_c_*, are calculated as follows:

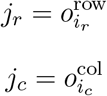

And the corresponding value in *M* is *M_jr,jc_*.

**InteractiveComplexHeatmap** has two functions `selectPosition()` and `selectArea()` which allow users to pick single positions or to select areas from heatmap. Under the interactive graphics device, users do not need to run `htPositionsOnDevice()` explicitly. The positions of heatmaps are automatically calculated, cached and reused if heatmaps are the same and the device has not changed its size. If users change the device size, `htPositionsOnDevice()` will be automatically re-executed.

Figure 8 demonstrates an example of using `selectPosition()`. Interactively, `selectPosition()` asks the user to click one position on the heatmap. The function returns a `DataFrame` which contains the heatmap name, slice name and row and column index in the corresponding matrix. An example output from `selectPosition()` is as follows:

**Figure 8:**
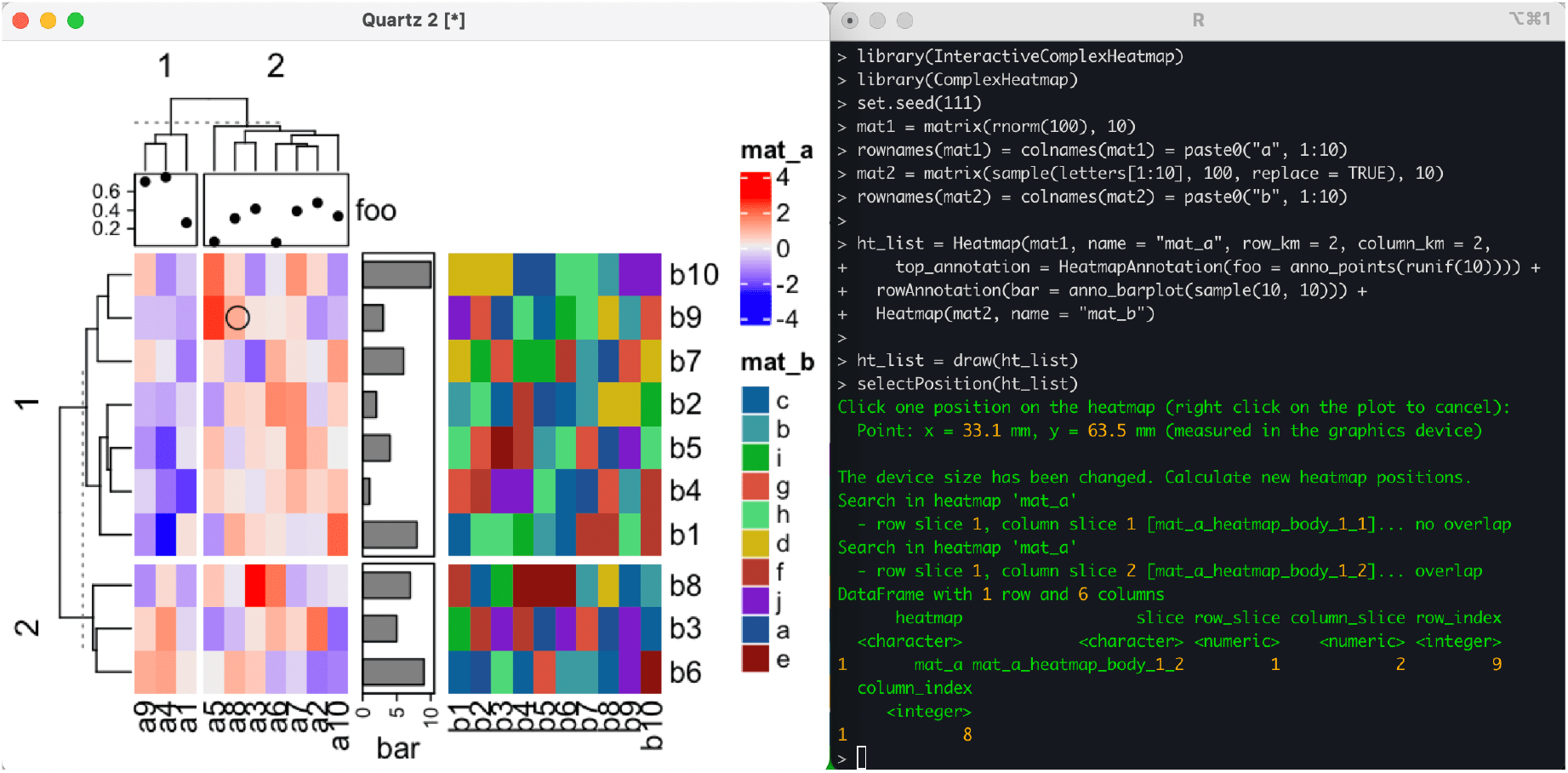
Demonstration of `selectPosition()`. Circle on the heatmap represents the point clicked by user. On the left is the interactive graphics device and on the right is the `R` terminal.

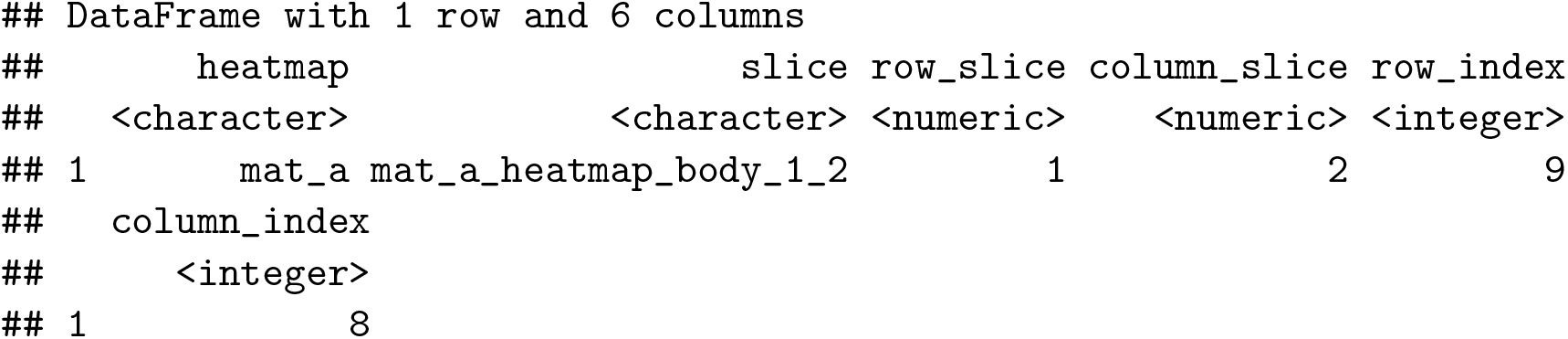

The output means the position that the user clicked is in a heatmap named `“mat_a”` and in its first row slice and in the second column slice. Assume `mat` is the matrix for heatmap `“mat_a”`, then the clicked point corresponds to the value `mat` [9, 8].

Similarly, the function `selectArea()` asks the user to click two positions on the heatmap which define an area. Note since the selected area may overlap over multiple heatmap slices, `selectArea()` returns a `DataFrame` with multiple rows. An example output is as follows.

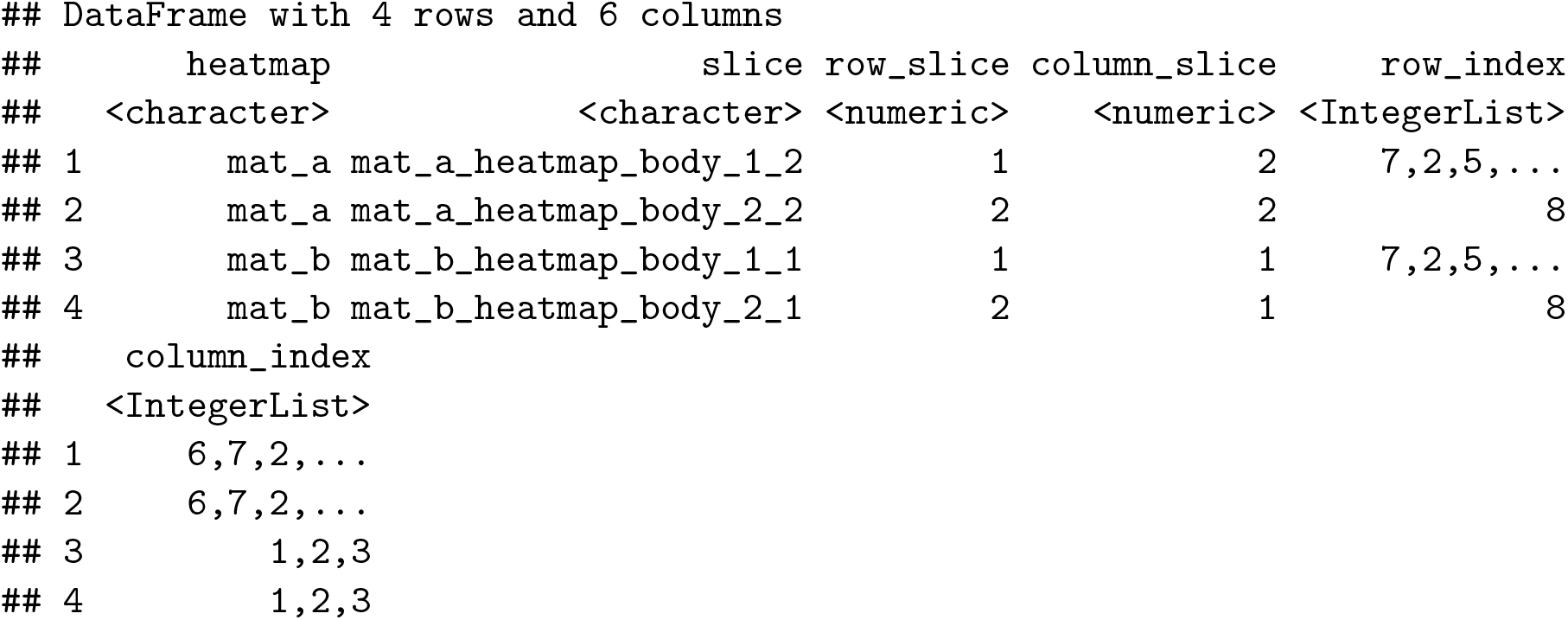

The columns `row_index` and `column_index` are stored in a class of `IntegerList` which is simply a list of integer vectors. To get the row indices in *e.g*. `mat_a_heatmap_body_1_2` (in the first row), user can use either one of the following commands:

~~~
*R> df[1, “row_index”][[1]] # assume the DataFrame object is called ‘df’
R> unlist(df[1, “row_index”])
R> df$row_index[[1]]*
~~~

### 4.2. On off-screen graphics devices

It is also possible to use `selectPosition()` and `selectArea()` on off-screen graphics devices, such as `pdf` or `png`. Now the positions cannot be selected interactively, but instead argument `pos` in `selectPosition()` or `pos1/pos2` in `selectArea()` need to be specified to simulate clicks. Values for `pos`, `pos1` and `pos2` should be `unit` objects of length two which corresponds to the *x* and *y* coordinates relative to the bottom left of the device.

~~~
*R> png(…)
R> ht_list = draw(ht_list)
R> pos = selectPosition(ht_list, pos = unit(c(3, 3), “cm”))
R> dev.off()*
~~~

Users do not need to use this functionality directly with an off-screen graphics device, but it is very useful when implementing the interactivity in a Shiny application where the plot is actually generated under an off-screen graphics device.

### 4.3. In Shiny web applications

With the three functions `htPositionsOnDevice()`, `selectPosition()` and `selectArea()`, it is possible to implement Shiny applications for interactively working with heatmaps. Now the question is how does the server side capture the positions that the user clicked on heatmap which is actually on the web page. Luckily, there is a solution. The heatmap is normally generated within a `shiny::plotOutput()` and `plotOutput()` provides two actions that can be applied on heatmaps on the web page: `click` and `brush`. Then under the Shiny framework, server can receive the information of the positions as soon as the user performs clicking or brushing on heamtaps. The positions can be later sent to `selectPosition()` or `selectArea()` via `pos` or `pos1/pos2` arguments for correctly corresponding to the original matrices. `htShiny()` is implemented in this way, as well as some other functions for Shiny application development, which will be introduced in the next section.

## 5. Shiny web application development

`htShiny()` exports heatmaps as stand-alone Shiny web applications. **InteractiveComplexHeatmap** also provides general solutions for integrating interactive heatmap widgets in general Shiny application development. It allows dynamically generating interactive heatmap widgets according to different configurations on heatmaps, and customizing the output that responds to user’s actions on heatmaps.

There are following two main functions for Shiny application development:

- `InteractiveComplexHeatmapOutput(),`
- `makelnteract iveComplexHeatmap().`

`InteractiveComplexHeatmapOutput()` constructs the user interface (UI). As already demonstrated in Section 3.2, the interactive heatmap widget by default contains three components, *i.e.*, the original heatmap, the sub-heatmap and the output that prints information of cells that users selected. The first two heatmap components contain various tools for controlling both heatmaps. `makeInteractiveComplexHeatmap()` defines a list of responses to user’s actions that were performed on heatmaps as well as in the tools.

Note `htShiny()` is simply a wrapper function on these two functions, thus, most of the functionalities introduced in this section also work for `htShiny()`.

### 5.1. With one interactive heatmap widget

Creating a Shiny application that only contains one interactive heatmap widget is rather simple. Following is an example that can be directly copied and pasted to an `R` session (Figure 9).

~~~
*R> library(ComplexHeatmap)
R> library(InteractiveComplexHeatmap)
R> library(shiny)
R>
R> data(rand_mat) # simply a random matrix
R> ht1 = Heatmap(rand_mat, name = “mat”,
+     show_row_names = FALSE, show_column_names = FALSE)
R> ht1 = draw(ht1)
R>
R> ui = fluidPage(
+     h3(“My first interactive ComplexHeatmap Shiny app”),
+     p(“This is an interactive heatmap visualization on a random matrix.”),
+     InteractiveComplexHeatmapOutput()
+)
R> server = function(input, output, session) {
+     makeInteractiveComplexHeatmap(input, output, session, ht1)
+  }
R> shinyApp(ui, server)*
~~~

**Figure 9:**
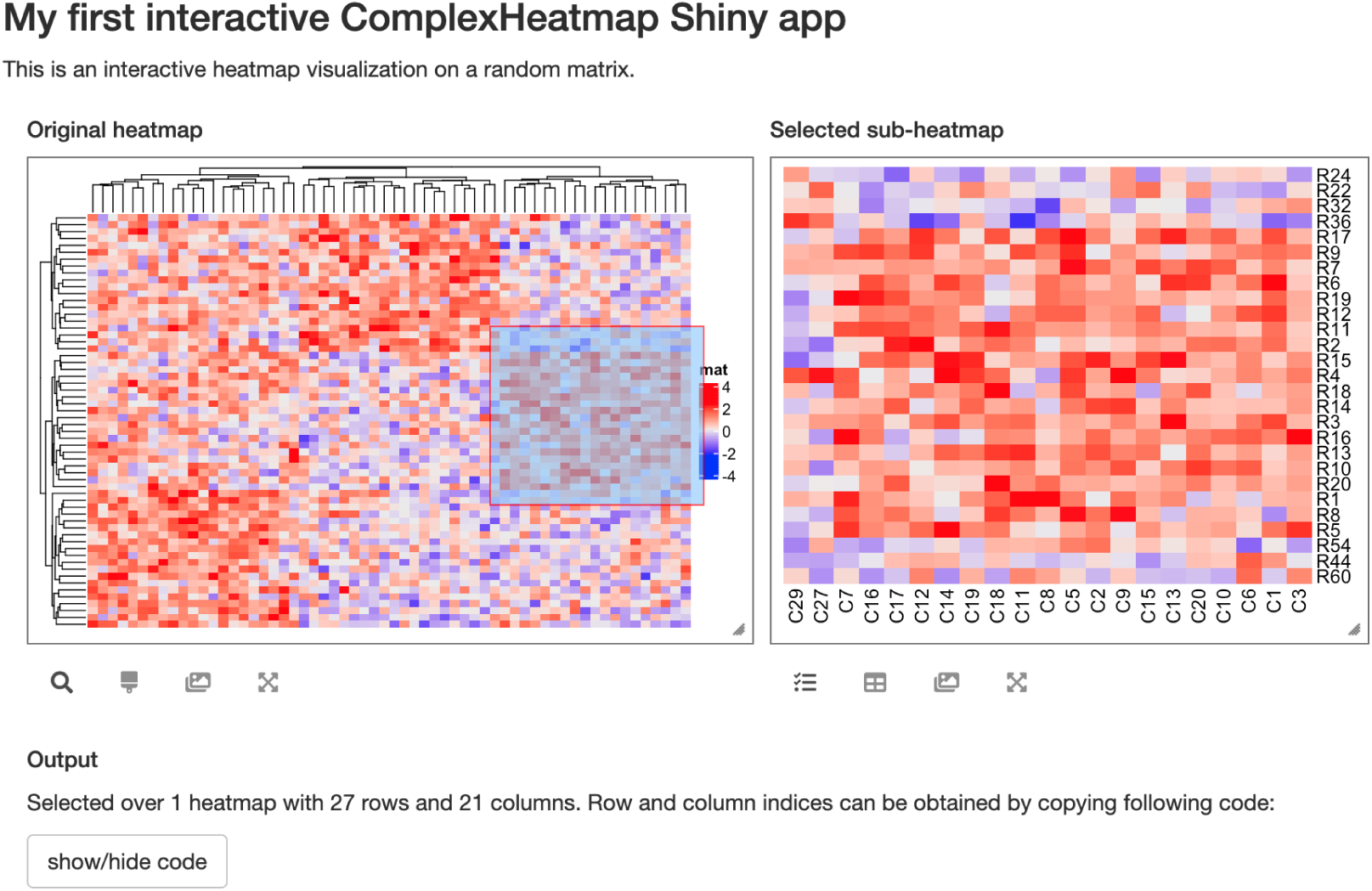
A simple Shiny application directly from ui and server.

### 5.2. With multiple interactive heatmap widgets

Multiple interactive heatmap widgets can be integrated into one single Shiny application. Now a “heatmap ID” must be assigned to each widget, so that `makeInteractiveComplexHeatmap()` can find the correct heatmap to respond. If there is only one heatmap widget in the Shiny application, such as in Section 5.1, the heatmap ID is internally generated if it is not specified and the UI is automatically linked to the server in the two functions. In the following example, we create a second widget which visualizes a random character matrix (Figure 10). The two interactive heatmap widgets are independent in the application. See `htShinyExample(1.8)` and `htShinyExample(5.1)` for live examples.

~~~
*R> set.seed(88)
R> mat2 = matrix(sample(letters[1:10], 100, replace = TRUE), 10)
R> ht2 = draw(Heatmap(mat2, name = “mat2”))
R>
R> ui = fluidPage(
+     h3(“The first heatmap”),
+     InteractiveComplexHeatmapOutput(“heatmap_1”,
+        height1 = 300, height2 = 300),
+     hr(),
+     h3(“The second heatmap”),
+     InteractiveComplexHeatmapOutput(“heatmap_2”,
+        height1 = 300, height2 = 300)
+)
R> server = function(input, output, session) {
+     makeInteractiveComplexHeatmap(input, output, session, ht1, “heatmap_1”)
+     makeInteractiveComplexHeatmap(input, output, session, ht2, “heatmap_2”)
+ }
R> shinyApp(ui, server)*
~~~

**Figure 10:**
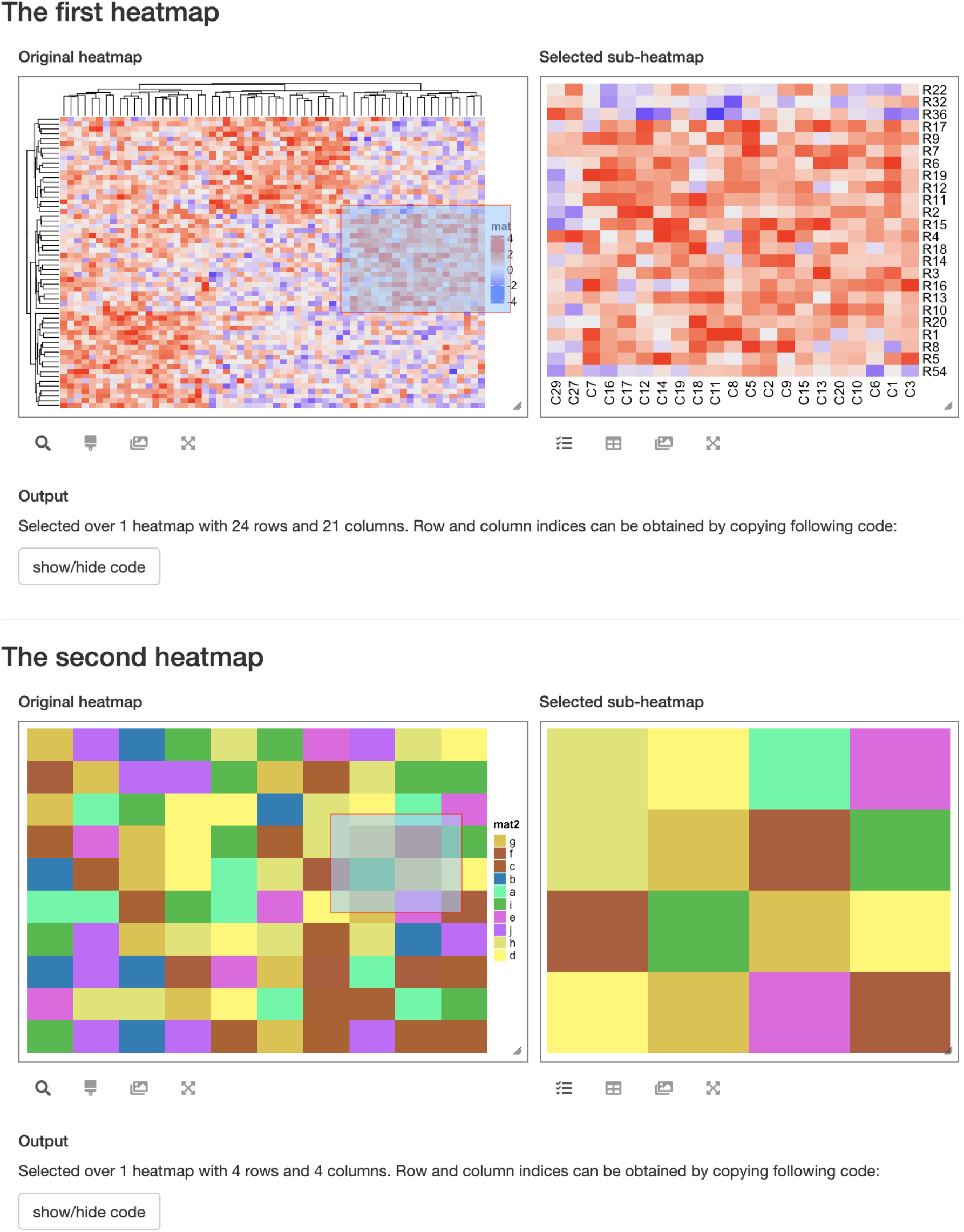
A Shiny application with two interactive heatmap widgets.

### 5.3. Customize the widgets

The original heatmap and sub-heatmap components can be resized by dragging from bottom right of the two boxes, but still, `InteractiveComplexHeatmapOutput()` provides arguments of `width1`, `height1`, `width2` and `height2` to precisely control the initial sizes of the two components. They can be manually set to make sure the heatmaps are well visualized and aligned when the application is initialized.

#### The layout

The layout of the three components is controlled by argument `layout` in `InteractiveComplexHeatmapOutput()`. It supports following values:

- `“(1-2)|3”`: Original heatmap and sub-heatmap are in the same row, and output is in the second row. This is the default layout.
- `“1|(2-3)”`: Original heatmap is in a single row, while sub-heatmap and output are in the second row.
- `“1-2-3”`: All three components are in the same row.
- `“1|2|3”`: Each component is in a single row.
- `“1-(2|3)”`: Original heatmap is in a single column. Sub-heatmap and output are vertically aligned and are in the second column. An example can be found with `htShinyExample(4.1)`.

Note the values for `layout` are in a special format to help to understand the layout, where the three code `1`, `2` and `3` correspond to original heatmap, sub-heatmap and output respectively, symbol `“-”` corresponds to horizontal alignment and `“|”` corresponds to vertical alignment. With different layouts, different default values are assigned to widths and heights of the three components to make sure they are well aligned.

#### Action on single heatmap cells

By default, to get the information of a single cell in the heatmap, a `“click”` action is used. In `InteractiveComplexHeatmapOutput()`, the action can also be set to `“hover”` or `“dblclick”`, then hovering or double clicking on heatmap will trigger the response on the sever side. The example in `htShinyExample(1.9)` demonstrates usages of these three actions.

#### Which action to respond

The argument `response` in `InteractiveComplexHeatmapOutput()` can be set as a vector with values in `“click”`, `“hover”`, `“dblclick”`, `“brush”` and `“brush-output”` to only respond to one or multiple events on heatmap. *E.g.*, if response is only set to `“click”`, there will be no response for the brush event in the interactive heatmap, also the sub-heatmap component is removed from the application.

Brushing on heatmap by default triggers two responses, one in the sub-heatmap and one in the output. If `“brush-output”` is included in `response` instead of `“brush”`, there will only be response in the output component, while the sub-heatmap component is removed from the application.

#### Separately specify the three UI components

`InteractiveComplexHeatmapOutput()` contains all three UI components. Nevertheless, the three components can be separately specified by three individual functions: `originalHeatmapOutput()`, `subHeatmapOutput()` and `HeatmapInfoOutput()`. This provides flexibility for the UI arrangement, e.g., to integrate with package **shinydashboard**(Chang and Borges Ribeiro 2018) where each UI component is wrapped within an individual box. An example is as follows (Figure 11).

~~~
*R> library(shinydashboard)
R> body = dashboardBody(
+     fluidRow(
+         box(title = “Original heatmap”, width = 4,
+             solidHeader = TRUE, status = “primary”,
+             originalHeatmapOutput(“ht”, title =NULL)),
+          box(title = “Sub-heatmap”, width = 4,
+             solidHeader = TRUE, status = “primary”,
+             subHeatmapOutput(“ht”, title = NULL)),
+         box(title = “Output”, width = 4,
+             solidHeader = TRUE, status = “primary”,
+             HeatmapInfoOutput(“ht”, title = NULL))
+)
+)
R> ui = dashboardPage(
+     dashboardHeader(),
+     dashboardSidebar(disable = TRUE),
+     body
+)
R> server = function(input, output, session) {
+     makeInteractiveComplexHeatmap(input, output, session, ht1, “ht”)
+ }
R> shinyApp(ui, server)*
~~~

**Figure 11:**
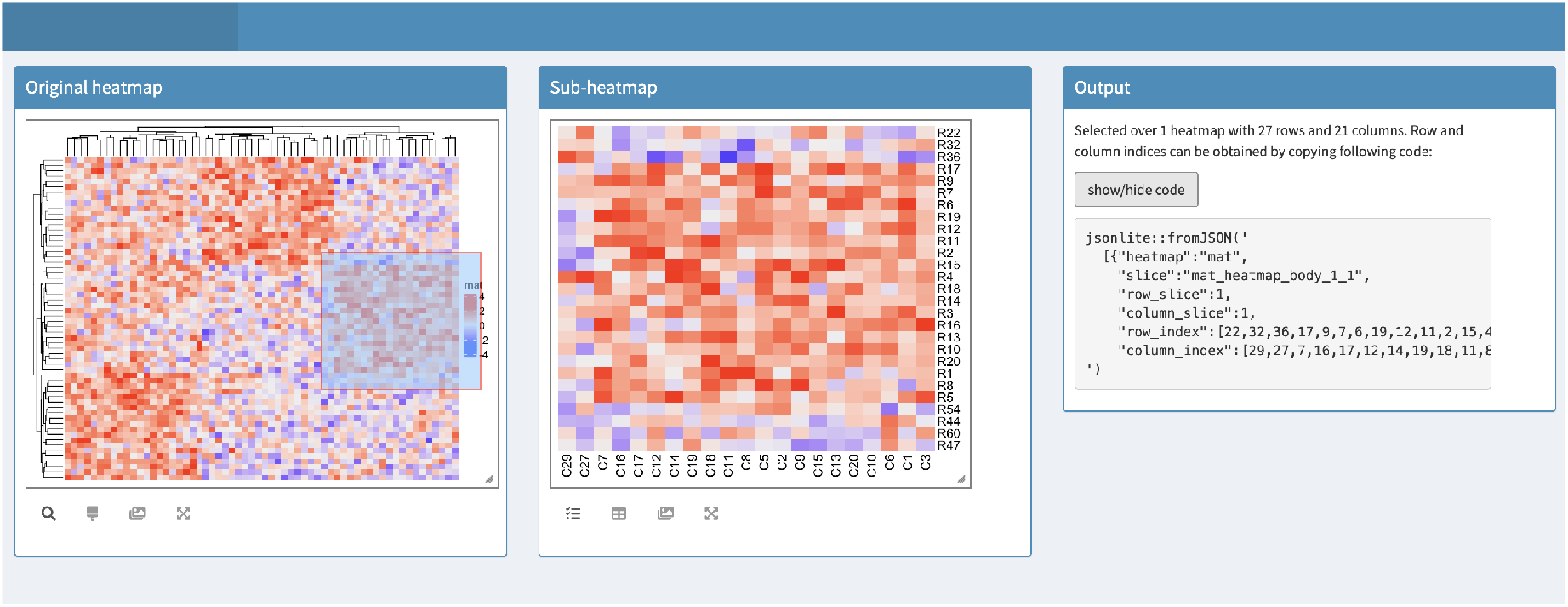
Separately specify the three UI components and integrate with **shinydashboard**.

Please note, since now the three components are generated independently, to correctly connect the three components as well as the server, the heatmap ID must be explicitly specified in all functions. More examples on integrating with **shinydashboard** can be found in `htShinyExample(10.1)` to `htShinyExample(10.5)`.

### 5.4. Work with R Markdown documents

It is straightforward to integrate **InteractiveComplexHeatmap** in an interactive R Markdown document, just in the same way as integrating normal Shiny applications. Examples can be found in `htShinyExample(7.1)` and `htShinyExample(7.2)`.

### 5.5. Self-define the output

Both clicking and brushing on a heatmap trigger an output associated with the heatmap. The output gives information of rows and columns selected by users. In **InteractiveComplexHeatmap**, the response for the two actions can be self-defined, which makes it possible to build an interactive application with complex responses.

There are two arguments: `click_action` and `brush_action` in `makeInteractiveComplex-Heatmap()` which accept self-defined functions to define how to respond after the heatmap was clicked or brushed. The two functions should accept two arguments, the first is a `DataFrame` object which contains the information of which rows and columns are selected by the user, and the second argument should always be `output` which is used in the Shiny server function. To use `click_action` or `brush_action`, a `htmlOutput` (or other similar *`Output`) should be first set up in the UI, then the Shiny application knows where to update the output. The output UI can replace the default output by directly assigning to argument `output_ui` in `InteractiveComplexHeatmapOutput()`.

~~~
*R> ui = fluidPage(
+     InteractiveComplexHeatmapOutput(output_ui = htmlūutput(“info”))
+)*
~~~

Or to create a new output UI independent to the interactive heatmap widget.

~~~
*R> ui = fluidPage(
+     InteractiveComplexHeatmapOutput(),
+     htmlOutput(“info”)
+)*
~~~

`click_action` or `brush_action` is normally defined as follows (assume the ID set in `htmlOutput()` is `“info”`):

~~~
*R> function(df, output) {
+     output$info = renderUI({
+         if(is.null(df)) {
+             …
+         } else {
+             …
+         }
+    })
+ }*
~~~

If users didn’t click or brush inside the heatmap body (*e.g*., only clicked in the dendrogram), `df` that is passed to the functions will be `NULL`. A sanity check might be performed here to make specific response when heatmap cell was not selected.

The format of `df` is slightly different between click and brush. If it is a click action, `df` has the same format as the returned object of `selectPosition()` (introduced in Section 4), which looks like the following chunk. It always has one row if the user clicked into the heatmap.

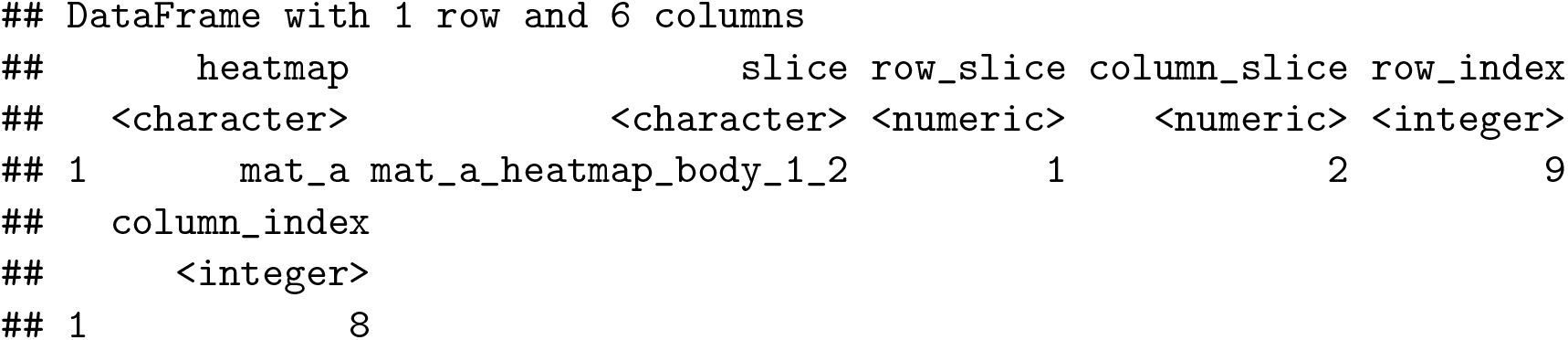

If it is a brush action, `df` has the same format as the returned object of `selectArea()` (introduced in Section 4), which looks like the following chunk. Each line contains row and column indices of the selected sub-matrix in a specific heatmap slice of a specific heatmap.

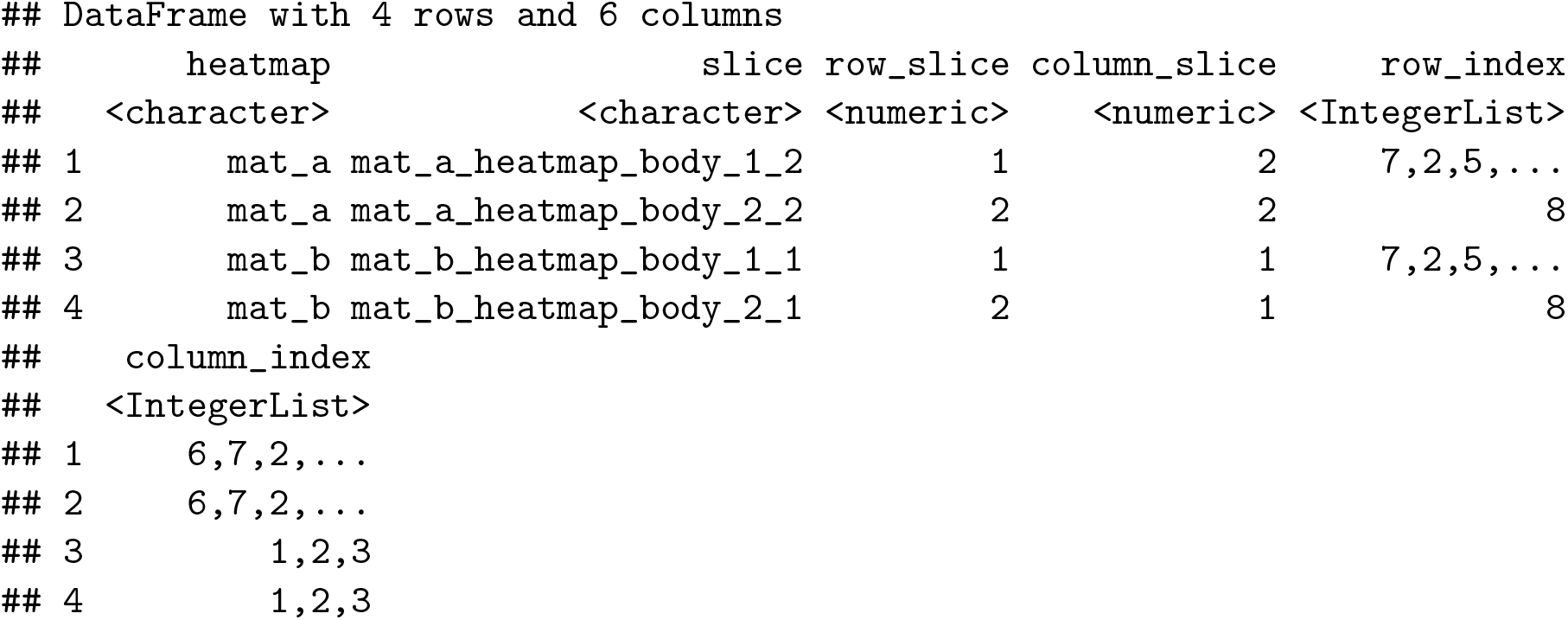

Note as demonstrated above, after a brush action, values in column `row_index` and `column_index` might be duplicated due to the fact that the selected heatmap slices are in the same row or column. *E.g*., in the previous example, the first and the third rows correspond to the selection in the first row slice, but in two different column slices, thus they have the same values for `row_index`. To safely get row indices and column indices of the selected heatmap, users might need to perform:

~~~
*R> unique(unlist(df$row_index))
R> unique(unlist(df$column_index))*
~~~

To receive more information from the Shiny application, the self-defined functions for `click_action` and `brush_action` can also accept four arguments where two additional variables `input` and session are passed from server function:

~~~
*R> function(df, input, output, session) {
+     …
+ }*
~~~

If argument `action` in `InteractiveComplexHeatmapOutput()` is set to `“hover”` or `“dblclick”`, the argument in `makeInteractiveComplexHeatmap()` is `hover_action` or `dblclick_action` correspondingly. Their usages are the same as `click_action`.

We demonstrate how to self-define `click_action` in the next example which visualizes a correlation matrix based on `mtcars` dataset. In the UI, the default output is replaced with a new `plotOutput`. On the server side, a self-defined `click_action` is defined to draw a scatter plot of the two corresponding variables that were selected from heatmap (Figure 12). Note in the example `response = “click”` is set to simplify Figure 12 so that the sub-heatmap component is not included in the application. The live version of this example can be found in `htShinyExample(5.5)`.

**Figure 12:**
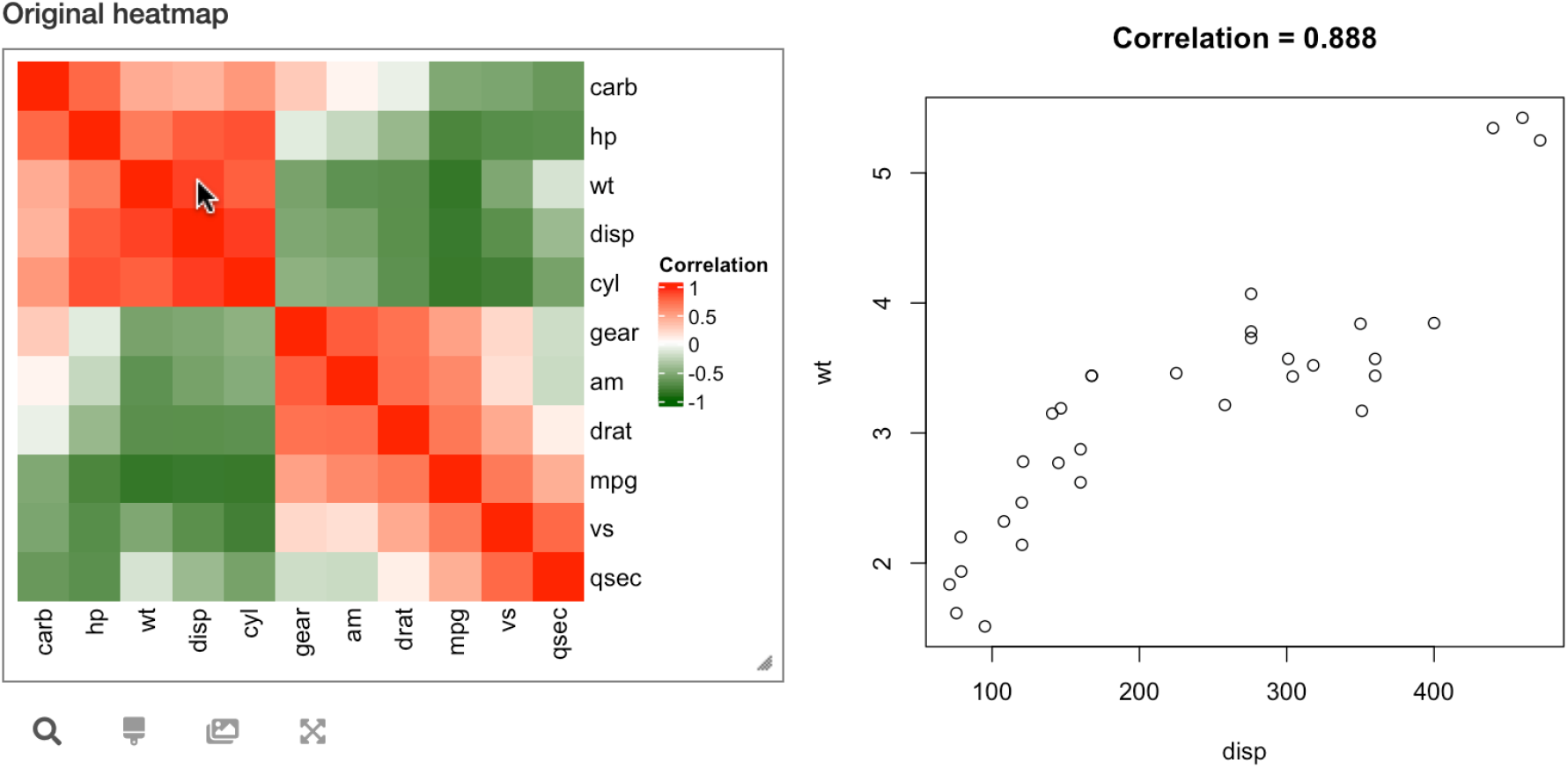
A demonstration of self-defining the output. The heatmap visualizes a correlation matrix based on **mtcars** dataset. Clicking on a cell draws a scatter plot of the two corresponding variables.

~~~
*R> data(mtcars)
R> cor_mat = cor(mtcars)
R>
R> # colorRamp2() from circlize is used to exactly define the color mapping
R> library(circlize)
R> col_fun = colorRamp2(c(-1, 0, 1), c(“darkgreen”, “white”, “red”))
R> ht = Heatmap(cor_mat, name = “Correlation”, col = col_fun,
+     show_row_dend = FALSE, show_column_dend = FALSE)
R>
R> ui = fluidPage(
+     InteractiveComplexHeatmapOutput(
+          output_ui = plotOutput(“scatterplot”, width = 400, height = 400),
+          response = “click”)
+)
R> click_action = function(df, output) {
+      output$scatterplot = renderPlot({
+          nm = colnames(mtcars)
+          i1 = df$column_index
+          i2 = df$row_index
+          x = mtcars[, i1]
+          y = mtcars[, i2]
+
+          plot(x, y, xlab = nm[i1], ylab = nm[i2],
+               main = paste0(“Correlation = ”, sprintf(‘%.3f’, cor(x, y))))
+    })
+ }
R> server = function(input, output, session) {
+     makeInteractiveComplexHeatmap(input, output, session, ht,
+         click_action = click_action)
+ }
R> shinyApp(ui, server)*
~~~

There are several other examples in **InteraciveComplexHeatmap** that demonstrate the use of self-defining output:

- `htShinyExample(5.3)`: A gene expression matrix is visualized and clicking on the heatmap prints the corresponding gene ID, the link to public database and other annotations related to this gene.
- `htShinyExample(5.4)`: The heatmap visualizes pairwise similarities of a list of Gene Ontology (GO) terms. In this example, the click and brush actions are self-defined so that the selected GO IDs as well as their detailed descriptions are printed.
- `htShinyExample(5.6)`: The heatmap visualizes pairwise Jaccard coefficients for multiple lists of genomic regions. Clicking on the heatmap cell draws a Hilbert curve generated by the package **HilbertCurve**(Gu, Eils, and Schlesner 2016b) which illustrates how the two corresponding sets of genomic regions overlap.
- Section 6 demonstrates a complex example of interactively visualizing results from a differential expression analysis where the output is self-defined to link selected genes on heatmaps to a MA-plot, a volcano plot as well as a table of results only for these selected genes.

#### Float the output

Instead of occupying static space in the application, the output component can be floated to the mouse positions by setting argument `output_ui_float = TRUE in InteractiveComplexHeatmapOutput()`, then clicking, hovering or brushing on the heatmap opens a floating frame that contains the output. `htShinyExample(9.1)` demonstrates output floating when mouse action is set to `“hover”`, `“click”` or `“dblclick”`, respectively. If the default output is replaced by a user-defined output by assigning to argument `output_ui`, the self-defined output can also be floated. Figure 13 demonstrates an example where clicking on the GO similarity heatmap opens a floating frame that contains the information of the corresponding GO terms. The live version can be found in `htShinyExample(9.2)`.

**Figure 13:**
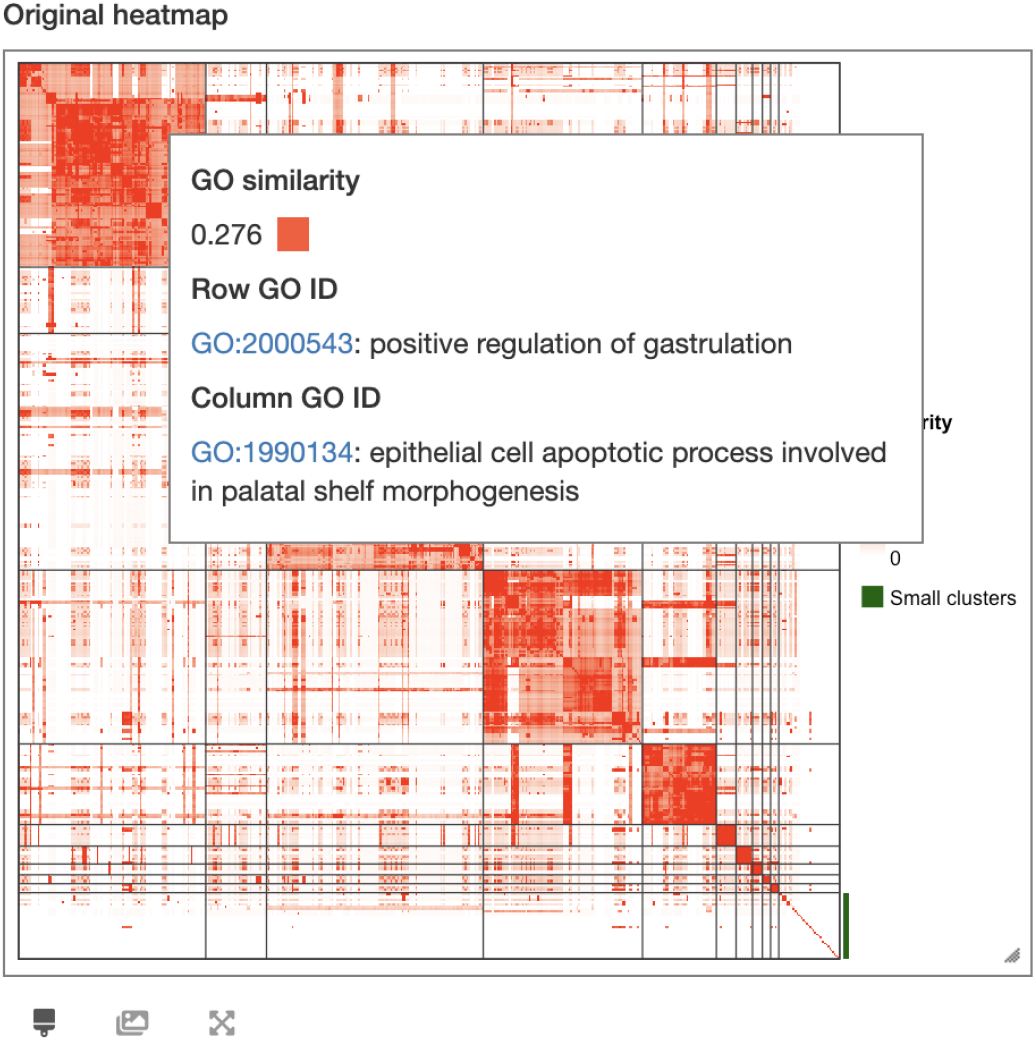
Floating output to mouse positions. The heatmap visualizes a GO similarity matrix from an analysis by package **simplifyEnrichment**. The output for clicking was self-defined to print the description of the two GO IDs and argument `output_ui_float in Interactive-ComplexHeatmapOutput()` was set to `TRUE` to float the output to mouse positions. Top left of the output box corresponds to the cell that was clicked.

#### Compact mode

`InteractiveComplexHeatmapOutput()` supports a “compact mode” by setting the argument compact to TRUE, then there is only the original heatmap and the output floats at the mouse positions if hovering or clicking on heatmap. The following two lines of code are atcually identical. An example is in `htShinyExample(1.11)`.

~~~
*R> InteractiveComplexHeatmapOutput(…, compact = TRUE)
R> InteractiveComplexHeatmapOutput(…, response = c(action, “brush-output”),
+    output_ui_float = TRUE)*
~~~

### 5.6. Dynamically generate interactive heatmap widget

In previous examples, heatmaps are already generated before generating the interactive applications. There are also scenarios where the heatmaps are generated on the fly. There might be following scenarios where heatmap is dynamically generated:

- The heatmap is based on a subset of matrix which is filtered by users, *e.g*., the expression matrix for differentially expressed genes filtered by different cutoffs.
- The heatmap annotations are dynamically provided by users.
- The heatmap parameters are selected by users, e.g., the clustering method or the splitting variable.
- If there are multiple heatmaps, which heatmaps are going to be drawn is dynamically selected.

**InteractiveComplexHeatmap** supports several ways to implement dynamic interactive heatmaps.

#### Directly use makeInteractiveComplexHeatmap()

The use is very natural. In the next example where the interactive heatmap widget is dynamically generated according to the number of heatmaps that the user selected, `makeInteractiveComplexHeatmap()` is directly put inside shiny: :`observeEvent()` or `shiny: :observe()` so that every time value of `input$n_heatmap` changes, it triggers an update of the interactive heatmap widget (Figure 14).

~~~
*R> ui = fluidPage(
+ sliderInput(“n_heatmap”, label = “How many heatmaps?”,
+     value = 1, min = 1, max = 5),
+ InteractiveComplexHeatmapOutput()
+)
R> generate_heatmap_list = function(n) {
+     ht_list = NULL
+     for(i in 1:n) {
+        ht_list = ht_list + Heatmap(matrix(rnorm(100), 10),
+           name = paste0(“mat_”, i),
+           column_title = paste0(“heatmap_”, i))
+    }
+    ht_list
+ }
R> server = function(input, output, session) {
+     observe({
+         ht_list = generate_heatmap_list(input$n_heatmap)
+         makeInteractiveComplexHeatmap(input, output, session, ht_list)
+  })
+ }
R> shiny::shinyApp(ui, server)*
~~~

**Figure 14:**
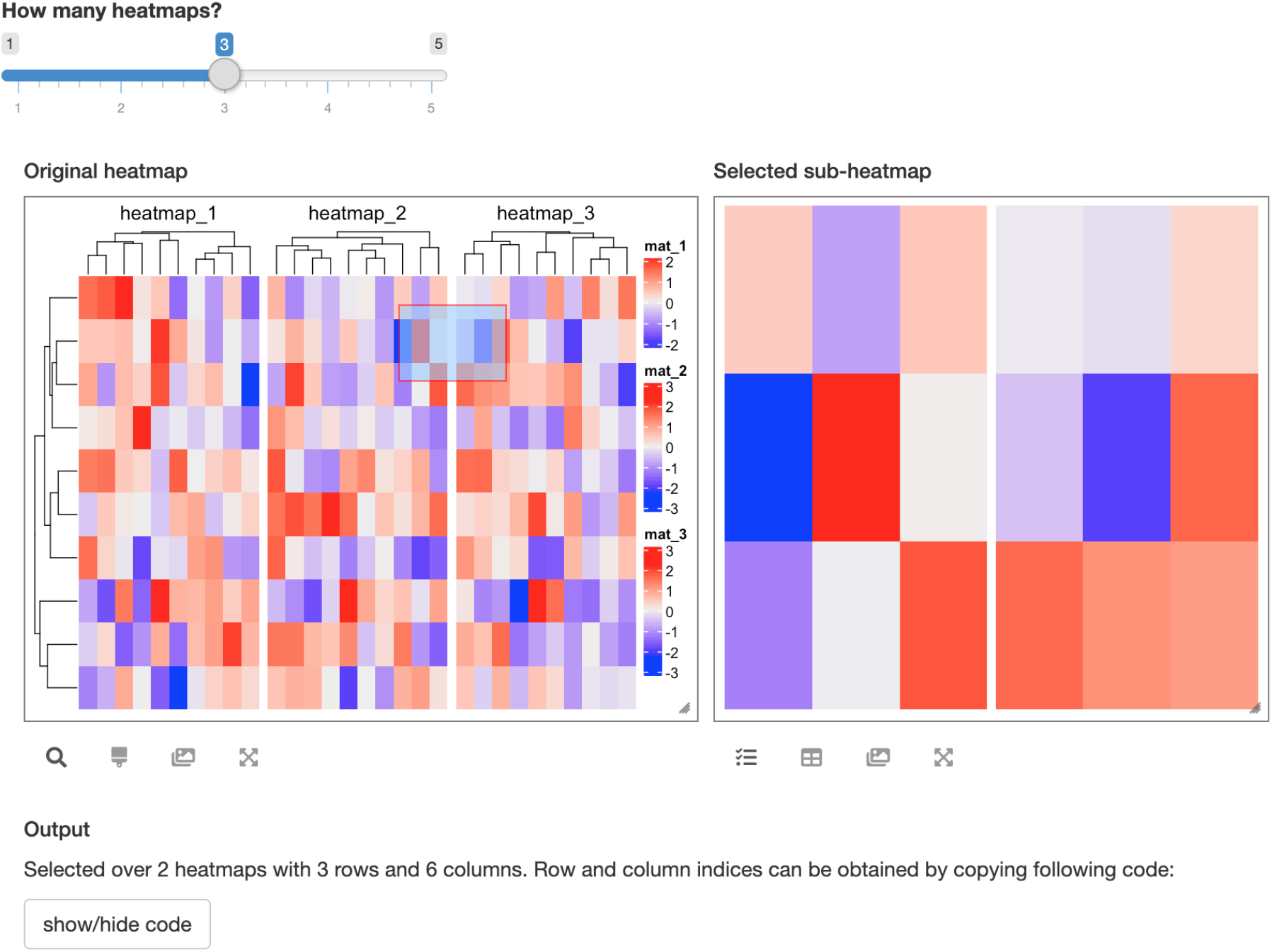
Dynamically generate the interactive heatmap widget. The wiget visualizes a user-specified number of heatmaps.

More examples can be found in `htShinyExample (6.1)` and `htShinyExample (6.2)`.

#### Use InteractiveComplexHeatmapWidget() and InteractiveComplexHeatmapModal()

In the previous example, the UI for the interactive heatmap must be manually created and it is placed on the application permanently. The function `InteractiveComplexHeatmapWidget()` provides an alternative way to implement this example which dynamically creates the complete interactive heatmap widget on the fly. To use `InteractiveComplexHeatmapWidget()`, a placeholder by `htmlOutput()` should be first created on the UI (e.g., `“heatmap_widget”` in the following code chunk). On the server side, `InteractiveComplexHeatmapWidget()` dynamically creates the UI and inserts it to the place defined by `htmlOutput()`. One advantage of using `InteractiveComplexHeatmapWidget()` is it supports removing the interactive heatmap widget from the application, which would be useful if the interactive widget is only for temporary use. See examples in `htShinyExample(6.6)` and `htShinyExample(6.7)`.

~~~
*R> ui = fluidPage(
+     sliderInput(“n_heatmap”, label = “How many heatmaps?”,
+         value = 1, min = 1, max = 5),
+     htmlOutput(“heatmap_widget”)
+)
R> server = function(input, output, session) {
+    observe({
+      ht_list = generate_heatmap_list(input$n_heatmap)
+      InteractiveComplexHeatmapWidget(input, output, session, ht_list,
+          output_id = “heatmap_widget”)
+   })
+ }
R> shiny::shinyApp(ui, server)*
~~~

The second function `InteractiveComplexHeatmapModal()` is very similar as `Interactive-ComplexHeatmapWidget()`. `InteractiveComplexHeatmapModal()` also dynamically generates the complete interactive heatmap widget on the fly, but it does not require to allocate a placeholder on UI, while the widget locates in a new layer above the current layer, as a so-called “modal dialog”.

Following code demonstrates an example which dynamically creates an interactive widget of a numeric heatmap or a character heatmap. Live examples can be found in `htShinyExample(6.3)`, `htShinyExample(6.4)` and `htShinyExample(6.5)`. The recursive interactive heatmap introduced in Section 3.4 is also implemented with `InteractiveComplexHeatmap-Modal()`.

~~~
*R> ui = fluidPage(
+     radioButtons(“select”, “Select a matrix:”,
+         choices = c(“Numeric” = 1, “Character” = 2)),
+     actionButton(“show_heatmap”, “Generate_heatmap”),
+)
R> server = function(input, output, session) {
+     observeEvent(input$show_heatmap, {
+     i = as.numeric(input$select)
+     if(i == 1) {
+          mat = matrix(rnorm(100), 10)
+      } else {
+          mat = matrix(sample(letters[1:10], 100, replace = TRUE), 10)
+      }
+      ht = Heatmap(mat)
+      InteractiveComplexHeatmapModal(input, output, session, ht)
+   })
+ }
R> shiny::shinyApp(ui, server)*
~~~

### 5.7. Implement the widgets from scratch

`InteractiveComplexHeatmapOutput()` and `makeInteractiveComplexHeatmap()` generate heatmap widgets that contain many pre-defined tools. **InteractiveComplexHeatmap** also provides low-level functions that directly return the information of rows and columns that were selected from heatmap so that users can define how to respond to the events on heatmap and build their own interactive heatmap widgets completely from scratch. We demonstrate the usage in the next example where `ui` and `server` are defined as follows:

~~~
*R> ui = fluidPage(
+     …,
+     plotOutput(“heatmap”, brush = “heatmap_brush”)
+)
R> server = function(input, output, session) {
+     ht_obj = reactiveVal(NULL)
+     ht_pos_obj = reactiveVal(NULL)
+
+ output$heatmap = renderPlot({
+    …
+    ht = draw(Heatmap(mat))
+    ht_pos = htPositionsOnDevice(ht)
+
+    ht_obj(ht)
+    ht_pos_obj(ht_pos)
+ })
+ observeEvent(input$heatmap_brush, {
+    pos = getPositionFromBrush(input$heatmap_brush)
+    df = selectArea(ht_obj (), pos[[1]], pos[[2]], mark = FALSE,
+          ht_pos = ht_pos_obj(), verbose = FALSE)
+
+    # the selected submatrix is captured in ‘df’,
+    # do something with ‘df’
+ …
+    })
+ }*
~~~

There are two points that need to be noted. 1. `draw()` and `htPositionsOnDevice()` should always be executed together and be put inside `renderPlot()` so that positions of all heatmap slices can be correctly calculated. 2. Use `getPositionFromBrush()` to retrieve the positions of the brushed area on heatmap, then the positions can be sent to `selectArea()` to correspond to the original matrix. Similarly, `getPositionFromClick()` and `selectPosition()` work together to retrieve row and column from the matrix that correspond to user’s click on the heatmap. Runnable examples can be found in `htShinyExample(5.7)`. This method is general that it also works for complex heatmaps, *e.g*., with row or column splitting, or with multiple heatmaps and annotations.

`htShinyExample(5.8)` demonstrates another example which visualizes a 2D density distribution. When brushing on the density heatmap, the use of `makeInteractiveComplexHeatmap()` which only purely visualizes a “zoomed subset” might cause a problem that is the densities in the selected sub-heatmap cannot be distinguished to background if densities are too small compared to the maximal density value from the complete dataset. To solve this problem, following the method proposed in this section, we reimplement the response of the brush event from scratch, which now triggers a new 2D density estimation only on the subset of the selected data. More description on this example can be found in Section 6.2.

## 6. Case studies

### 6.1. Visualize differential gene expression results

Figure 15 demonstrates an interactive heatmap application which visualizes results of differential gene expression analysis on the **airway** dataset performed by package **DESeq2** (Love, Huber, and Anders 2014). In the application, the heatmap visualizes expression of genes that are significantly different from a two-condition comparison, *i.e.,* `trt` vs `untrt` (treated vs. untreated). The application is arranged in a three-column layout where the original heatmap locates in the first column, the sub-heatmap and the default output locate in the second column, and self-defined outputs are in the third column. The heatmap components are specified separately and the layout is implemented with the **shinydashboard** package.

**Figure 15:**
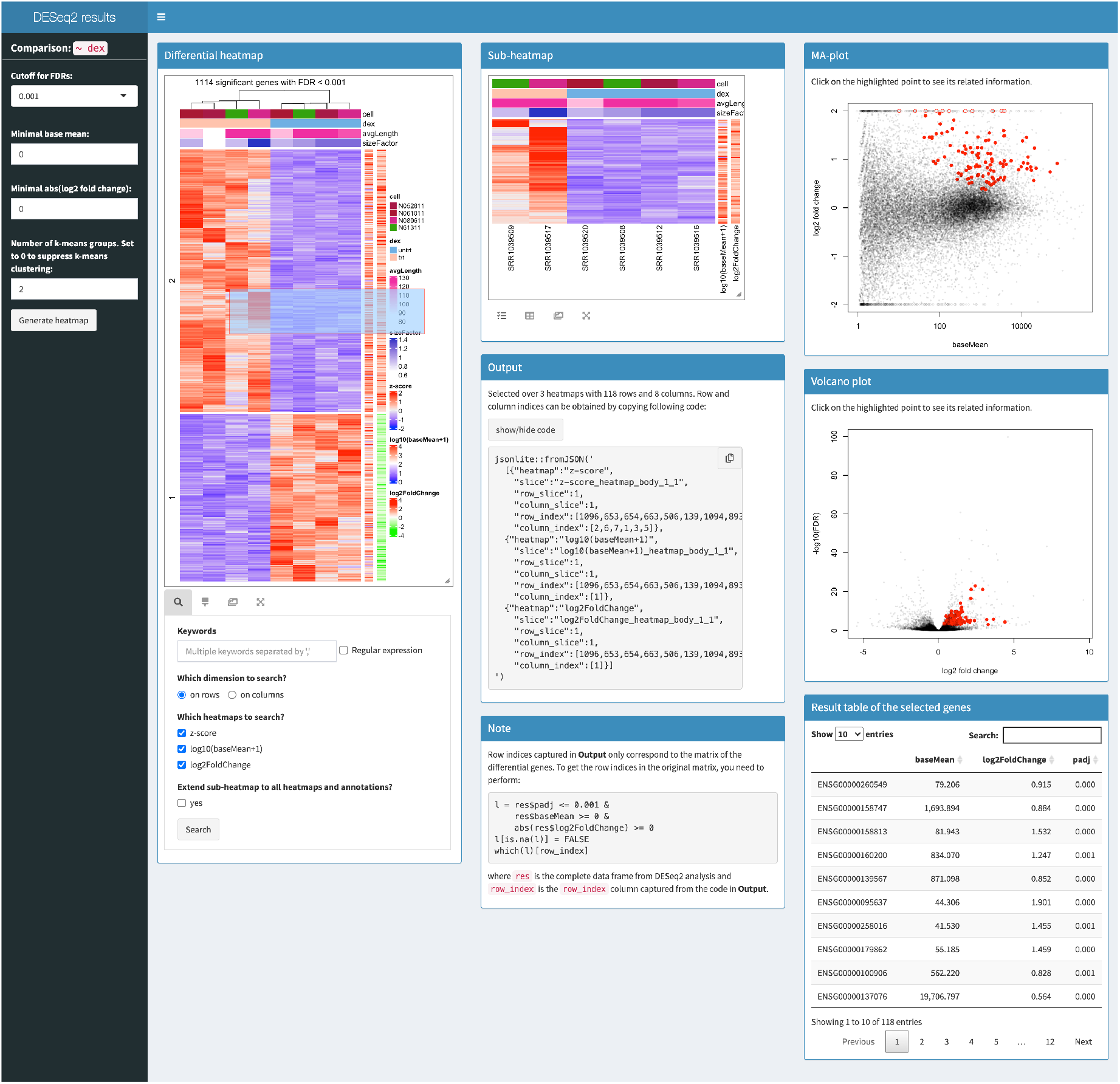
An interactive heatmap application for visualizing results from a differential gene expression analysis.

In the original heatmap list, there are two additional one-column heatmaps which visualize the absolute expression level of genes (`baseMean`) and the fold change between the two conditions (`log2foldChange`). To get more comprehensive information of how genes are differentially expressed when selecting them from heatmap, we defined extra outputs for the click and brush events on the original heatmaps. As shown in Figure 15, when a subset of genes are selected from the original heatmap, they are highlighted in a MA-plot which is a scatter plot of `log2foldChange` against `baseMean` where `baseMean` is in log10 scale and in a volcano plot which is also a scatter plot of `-log10(FDR)` against `log2foldchange`. There is a table of statistics from **DESeq2** analysis for the selected genes is printed below the two plots. Users can also search in heatmaps to obtain a subset of genes of interest to generate corresponding MA-plot, volcano plot and table.

In the left sidebar of the application, there are parameters for selecting differential genes as well as for selecting the number of groups for the *k*-means clustering applied on the expression matrix. The whole interactive widget will be updated when specific parameters are selected.

The vignette “A Shiny app for visualizing DESeq2 results” in **InteractiveComplexHeatmap** gives a detailed explanation of implementing this Shiny application from scratch. Because differential expression analysis with **DESeq2** is widely applied in transcriptomic data analysis, **InteractiveComplexHeatmap** implements a generic function `interactivate()` and a method dispatched on `DESeqDataSet` class which is defined by **DESeq2** for storing results from differential expression analysis. Thus, interactive visualization on **DESeq2** results can be simply achieved such as in the following example:

~~~
*R> library(DESeq2)
R> library(airway)
R> data(airway)
R>
R> dds = DESeqDataSet(airway, design = ~ dex)
R> dds = DESeq(dds)
R> interactivate(dds)*
~~~

### 6.2. Visualize two-dimensional density distributions

Heatmap is a popular method for visualizing two-dimensional density distributions. Nevertheless, when there are peaks that have large heights in the distribution, other peaks with smaller heights might be difficult to be distinguished from each other and even not distinguishable from background. Figure 16 implements an approach where selecting a region from the original density heatmap triggers a new two-dimensional density estimation but only on the selected subset of data. This approach helps to reveal more details in the regions which have relatively lower densities and are normally hidden in the complete dataset. To facilitate users to easier interprate two-dimensional distributions, **InteractiveComplexHeatmap** provides a general-purpose function `interactivateDensity2D()` that can be applied to two numeric vectors where the density is estimated by the package **ks** (Duong 2007).

~~~
*R> interactivateDensity2D(x, y)*
~~~

If the two-dimensional density is already estimated by function `ks: :kde()`, the generic function interactivate() can be directly applied.

~~~
*R> fit = ks::kde(…)
R> interactivate(fit)*
~~~

**Figure 16:**
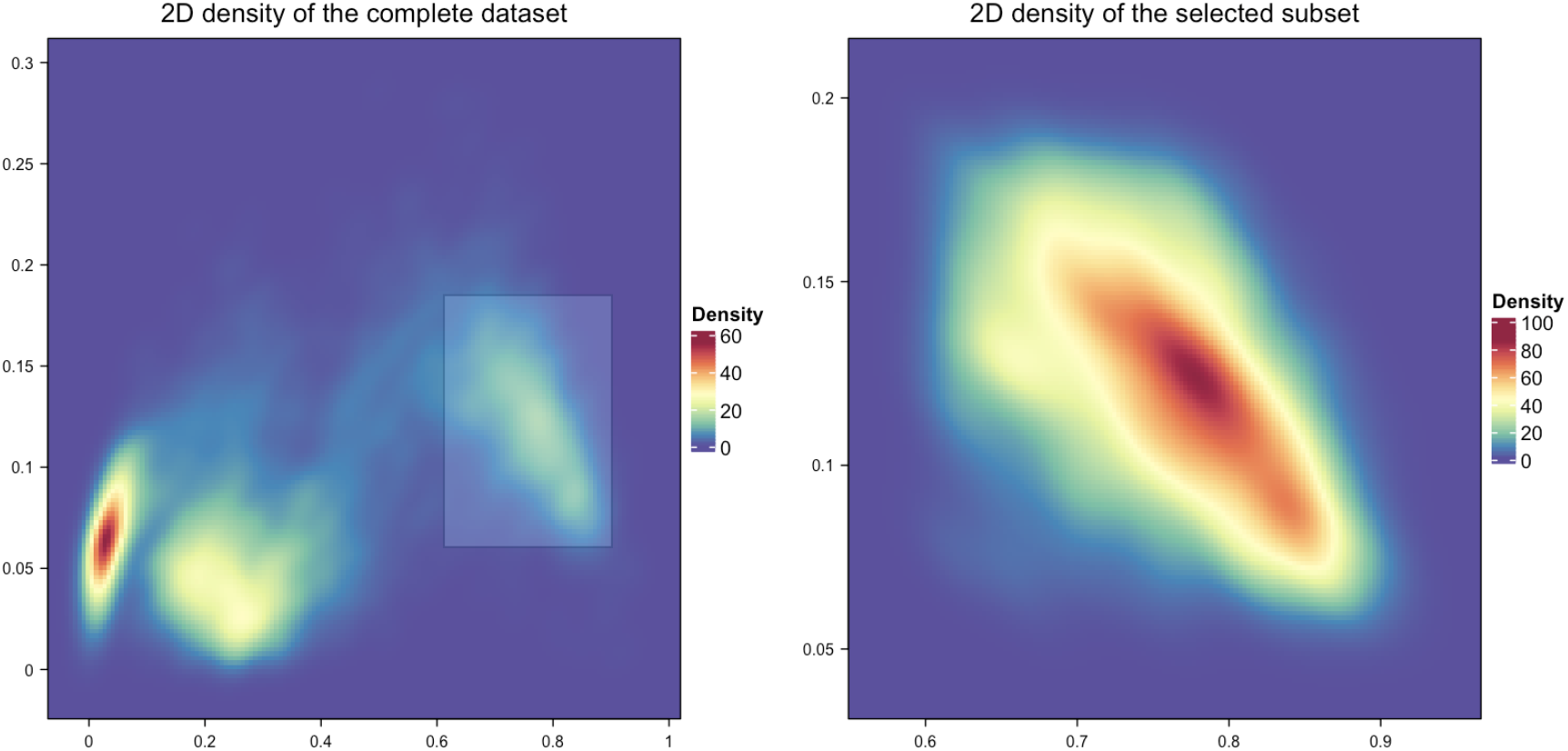
An interacitve visualization on a two-dimensional density distribution.

## 7. Conclusion

Interactivity on heatmaps greatly facilitates users to capture features which have specific patterns on heatmaps. In this paper, we described a new **`R`**/Bioconductor package **InteractiveComplexHeatmap** that supports interactive visualization on complex heatmaps. It can easily export static heatmaps to interactive Shiny applications and it also provides flexible functionalities for implementing more comprehensive Shiny applications. We believe it will be a useful tool for effectively interpreting data and developmenting new tools.

## 8. Acknowledgments

This work was supported by the German Cancer Research Center-Heidelberg Center for Personalized Oncology (DKFZ-HIPO) and by the Molecular Diagnostics Program of the National Center for Tumor Diseases (NCT) Heidelberg.

