## Supplementary file 1 for "Make Interactive Complex Heatmaps in R": InteractiveComplexHeatmap_manuscript_code.html

Code for generating the figures in manuscript ‘Make Interactive Complex Heatmaps in R’


### Code for generating the figures in manuscript ‘Make Interactive Complex Heatmaps in R’

###### Zuguang Gu

#### 2021-04-02

#### Figure 1

##### Figure 1A

```
library(ComplexHeatmap)
library(circlize)
library(GetoptLong)

set.seed(123)
nr1 = 4; nr2 = 8; nr3 = 6; nr = nr1 + nr2 + nr3
nc1 = 6; nc2 = 8; nc3 = 10; nc = nc1 + nc2 + nc3
mat = cbind(rbind(matrix(rnorm(nr1*nc1, mean = 1,   sd = 0.5), nr = nr1),
          matrix(rnorm(nr2*nc1, mean = 0,   sd = 0.5), nr = nr2),
          matrix(rnorm(nr3*nc1, mean = 0,   sd = 0.5), nr = nr3)),
    rbind(matrix(rnorm(nr1*nc2, mean = 0,   sd = 0.5), nr = nr1),
          matrix(rnorm(nr2*nc2, mean = 1,   sd = 0.5), nr = nr2),
          matrix(rnorm(nr3*nc2, mean = 0,   sd = 0.5), nr = nr3)),
    rbind(matrix(rnorm(nr1*nc3, mean = 0.5, sd = 0.5), nr = nr1),
          matrix(rnorm(nr2*nc3, mean = 0.5, sd = 0.5), nr = nr2),
          matrix(rnorm(nr3*nc3, mean = 1,   sd = 0.5), nr = nr3))
   )
mat = mat[sample(nr, nr), sample(nc, nc)] # random shuffle rows and columns
rownames(mat) = paste0("row", seq_len(nr))
colnames(mat) = paste0("column", seq_len(nc))

library(dendextend)
column_dend = as.dendrogram(hclust(dist(t(mat))))
column_dend = color_branches(column_dend, k = 3) # `color_branches()` returns a dendrogram object
column_dend = dendrapply(column_dend, function(d) {
    if(runif(1) > 0.5) attr(d, "nodePar") = list(cex = 0.8, pch = sample(20, 1), col = rand_color(1))
    return(d)
})
ht = Heatmap(mat, name = "mat", row_dend_width = unit(2, "cm"), cluster_columns = column_dend,
    column_names_rot = 45, row_split = rep(c("A", "B"), 9), row_km = 2, column_split = 3,
    top_annotation = HeatmapAnnotation(foo1 = 1:24, bar1 = anno_points(runif(24))),
    right_annotation = rowAnnotation(foo2 = 18:1, bar2 = anno_barplot(cbind(runif(18), runif(18)), 
        gp = gpar(fill = 2:3), width = unit(2, "cm"))))

draw(ht, annotation_legend_list = list(Legend(title = "bar2", labels = c("group1", "group2"), 
    legend_gp = gpar(fill = 2:3))),
    merge_legend = TRUE, padding = unit(c(5, 5, 5, 5), "mm"))
for(slice in 1:3) {
    decorate_annotation("bar1", slice = slice, {
        grid.lines(c(0, 1), c(0.5, 0.5), gp = gpar(lty = 2, col = "#AAAAAA"))
    })
}
```

##### Figure 1B

```
small_mat = mat[1:9, 1:9]
col_fun = colorRamp2(c(-2, 0, 2), c("green", "white", "red"))
ht = Heatmap(small_mat, name = "mat", col = col_fun,
    row_km = 2, column_km = 2,
    layer_fun = function(j, i, x, y, w, h, fill, slice_r, slice_c) {
        # restore_matrix() is explained after this chunk of code
        ind_mat = restore_matrix(j, i, x, y)
        for(ir in seq_len(nrow(ind_mat))) {
            # start from the second column
            for(ic in seq_len(ncol(ind_mat))[-1]) {
                ind1 = ind_mat[ir, ic-1] # previous column
                ind2 = ind_mat[ir, ic]   # current column
                v1 = small_mat[i[ind1], j[ind1]]
                v2 = small_mat[i[ind2], j[ind2]]
                if(v1 * v2 > 0) { # if they have the same sign
                    col = ifelse(v1 > 0, "violet", "blue")
                    grid.segments(x[ind1], y[ind1], x[ind2], y[ind2],
                        gp = gpar(col = col, lwd = 2))
                    grid.points(x[c(ind1, ind2)], y[c(ind1, ind2)], 
                        pch = 16, gp = gpar(col = col), size = unit(4, "mm"))
                }
            }
        }
        if(slice_r != slice_c) {
            grid.rect(gp = gpar(lwd = 2, fill = "transparent"))
        }
    }
)
draw(ht, padding = unit(c(5, 5, 5, 5), "mm"))
```

##### Figure 1C

```
image_png = sample(dir("~/project/development/ComplexHeatmap-reference/IcoMoon-Free-master/PNG/64px", 
    full.names = TRUE), 20)

set.seed(123)
n = 7
m = matrix(rnorm(100*n), nc = 100)
m2 = matrix(rnorm(100*n), nc = 100)

ha1 = rowAnnotation(
    "anno_simple" = runif(n),
    "anno_image" = anno_image(image_png[1:n], gp = gpar(col = "black"), 
        space= unit(2, "mm"), width = unit(1, "cm")),
    "anno_points" = anno_points(matrix(runif(2*n), nc = 2), pch = 1:2, 
        gp = gpar(col = 2:3), width = unit(2, "cm")),
    "anno_lines" = anno_lines(cbind(runif(n), runif(n)), 
        gp = gpar(col = 2:3, lty = 1:2), width = unit(2, "cm")),
    "anno_barplot" = anno_barplot(cbind(runif(n), runif(n)), 
        gp = gpar(fill = 2:3, col = 2:3), width = unit(2, "cm")),
    "anno_boxplot" = anno_boxplot(m, gp = gpar(fill = 1:n), 
        width = unit(2, "cm")),
    "anno_text" = anno_text(gt_render(qq("row<sub>@{1:n}</sub>", collapse = FALSE), 
                        align_widths = TRUE, halign = 0.5, 
                        r = unit(4, "pt"),
                        padding = unit(c(2, 8, 2, 8), "pt")), 
                    gp = gpar(box_col = "blue", box_lwd = 2, col = 1:10, fontfamily = "Times"), 
                    just = "right", 
                    location = unit(1, "npc")
    ),
    "anno_histogram" = anno_histogram(m2, gp = gpar(fill = 1:n), border = TRUE, 
        width = unit(3.5, "cm")),
    "anon_violin" = anno_density(m, type = "violin", 
        gp = gpar(fill = 1:10, col = c("white", rep("black", 7), "white", "black")), 
        width = unit(3.5, "cm")),
    show_legend = FALSE, simple_anno_size = unit(1, "cm"), gap = unit(2, "mm"), 
    show_annotation_name = FALSE
)


m = matrix(rnorm(1500), nc = 15)
lt1 = apply(m, 2, function(x) data.frame(density(x)[c("x", "y")]))

lt2 = lapply(1:15, function(x) cumprod(1 + runif(1000, -x/100, x/100)) - 1)

# 20 rows
ha3 = rowAnnotation(
    "anno_joyplot" = anno_joyplot(lt1, width = unit(4, "cm"), gp = gpar(fill = 1:10), 
        transparency = 0.75, scale = 2),
    "anno_horizon" = anno_horizon(lt2),
    gap = unit(2, "mm"), show_annotation_name = FALSE
)

grid.newpage()
grid.rect(gp = gpar(fill = "white", col = NA))
pushViewport(viewport(width = ComplexHeatmap:::width(ha1) + ComplexHeatmap:::width(ha3)))

pushViewport(viewport(x = 0, y = unit(1, "npc") - unit(5, "mm"), 
    width = ComplexHeatmap:::width(ha1), height = unit(1, "npc") - unit(2, "cm"), 
    just = c("left", "top")))
draw(ha1)
popViewport()

pushViewport(viewport(x = ComplexHeatmap:::width(ha1) + unit(2, "mm"), 
    y = unit(1, "npc") - unit(5, "mm"), 
    width = ComplexHeatmap:::width(ha3) - unit(2, "mm"), height = unit(1, "npc") - unit(2, "cm"), 
    just = c("left", "top")))
draw(ha3)
popViewport()

popViewport()
```

##### Figure 1D

```
res_list = readRDS(url("http://jokergoo.github.io/supplementary/ComplexHeatmap-supplementary1-4/supplS3_methylation/meth.rds"))
type = res_list$type
mat_meth = res_list$mat_meth
mat_expr = res_list$mat_expr
direction = res_list$direction
cor_pvalue = res_list$cor_pvalue
gene_type = res_list$gene_type
anno_gene = res_list$anno_gene
dist = res_list$dist
anno_enhancer = res_list$anno_enhancer

column_tree = hclust(dist(t(mat_meth)))
column_order = column_tree$order

library(RColorBrewer)
meth_col_fun = colorRamp2(c(0, 0.5, 1), c("blue", "white", "red"))
direction_col = c("hyper" = "red", "hypo" = "blue")
expr_col_fun = colorRamp2(c(-2, 0, 2), c("green", "white", "red"))
pvalue_col_fun = colorRamp2(c(0, 2, 4), c("white", "white", "red"))
gene_type_col = structure(brewer.pal(length(unique(gene_type)), "Set3"), 
    names = unique(gene_type))
anno_gene_col = structure(brewer.pal(length(unique(anno_gene)), "Set1"), 
    names = unique(anno_gene))
dist_col_fun = colorRamp2(c(0, 10000), c("black", "white"))
enhancer_col_fun = colorRamp2(c(0, 1), c("white", "orange"))

ht_opt(
    legend_title_gp = gpar(fontsize = 8, fontface = "bold"), 
    legend_labels_gp = gpar(fontsize = 8), 
    heatmap_column_names_gp = gpar(fontsize = 8),
    heatmap_column_title_gp = gpar(fontsize = 10),
    heatmap_row_title_gp = gpar(fontsize = 8)
)

ha = HeatmapAnnotation(type = type, 
    col = list(type = c("Tumor" = "pink", "Control" = "royalblue")),
    annotation_name_side = "left")
ha2 = HeatmapAnnotation(type = type, 
    col = list(type = c("Tumor" = "pink", "Control" = "royalblue")), 
    show_legend = FALSE)

ht_list = Heatmap(mat_meth, name = "methylation", col = meth_col_fun,
    column_order= column_order,
    top_annotation = ha, column_title = "Methylation") +
    Heatmap(direction, name = "direction", col = direction_col) +
    Heatmap(mat_expr[, column_tree$order], name = "expression", 
        col = expr_col_fun, 
        column_order = column_order, 
        top_annotation = ha2, column_title = "Expression") +
    Heatmap(cor_pvalue, name = "cor_p", col = pvalue_col_fun) +
    Heatmap(gene_type, name = "gene type", col = gene_type_col) +
    Heatmap(anno_gene, name = "anno_gene", col = anno_gene_col) +
    Heatmap(dist, name = "dist_tss", col = dist_col_fun) +
    Heatmap(anno_enhancer, name = "anno_enhancer", col = enhancer_col_fun, 
        cluster_columns = FALSE, column_title = "Enhancer")

draw(ht_list, row_km = 2, row_split = direction,
    column_title = "Comprehensive correspondence between methylation, expression and other genomic features", 
    column_title_gp = gpar(fontsize = 12, fontface = "bold"), 
    merge_legends = TRUE, padding = unit(c(5, 15, 5, 15), "mm"))
```

```
ht_opt(RESET = TRUE)
```

#### Figure 2

```
library(ComplexHeatmap)
set.seed(111)
mat1 = matrix(rnorm(100), 10)
rownames(mat1) = colnames(mat1) = paste0("a", 1:10)
mat2 = matrix(sample(letters[1:10], 100, replace = TRUE), 10)
rownames(mat2) = colnames(mat2) = paste0("b", 1:10)

ht_list = Heatmap(mat1, name = "mat_a", row_km = 2, column_km = 2,
        top_annotation = HeatmapAnnotation(foo = anno_points(runif(10)))) +
    rowAnnotation(bar = anno_barplot(sample(10, 10))) +
    Heatmap(mat2, name = "mat_b")

ht_list = draw(ht_list) # not necessary, but recommended

library(InteractiveComplexHeatmap)
htShiny(ht_list)
```

#### Figure 3

```
library(InteractiveComplexHeatmap)
library(cola)  # from Bioconductor
data(golub_cola)
get_signatures(golub_cola["ATC:skmeans"], k = 3) # this makes the heatmap
ht_shiny()
```

#### Figure 4

Following code opens a Shiny application which contains the souce code that generate it. It is the same for the following part of this document where `htShinyExample()` is used. We won’t repeatedly mention there.

```
htShinyExample(4.2)
```

#### Figure 5

##### Figure 5A

```
htShinyExample(2.5)
```

##### Figure 5B

```
htShinyExample(2.6)
```

##### Figure 5C

```
htShinyExample(2.4)
```

#### Figure 6

##### Figure 6A

```
htShinyExample(2.2)
```

##### Figure 6B

```
htShinyExample(2.3)
```

##### Figure 6C

```
htShinyExample(2.1)
```

##### Figure 6D

```
htShinyExample(3.1)
```

#### Figure 7

```
library(InteractiveComplexHeatmap)
library(ComplexHeatmap)
set.seed(111)
mat1 = matrix(rnorm(100), 10)
rownames(mat1) = colnames(mat1) = paste0("a", 1:10)
mat2 = matrix(sample(letters[1:10], 100, replace = TRUE), 10)
rownames(mat2) = colnames(mat2) = paste0("b", 1:10)

ht_list = Heatmap(mat1, name = "mat_a", row_km = 2, column_km = 2,
        top_annotation = HeatmapAnnotation(foo = anno_points(runif(10)))) +
    rowAnnotation(bar = anno_barplot(sample(10, 10))) +
    Heatmap(mat2, name = "mat_b")


library(cowplot)

p1 = grid.grabExpr({
    ht_list = draw(ht_list)
    pos = htPositionsOnDevice(ht_list)
    seekViewport("global")
    grid.rect(gp = gpar(lty = 2, fill = "transparent"))
}, width = 6, height = 4)

p2 = grid.grabExpr({
    grid.newpage()
    grid.rect(gp = gpar(lty = 2))
    for(i in seq_len(nrow(pos))) {
        x_min = pos[i, "x_min"]
        x_max = pos[i, "x_max"]
        y_min = pos[i, "y_min"]
        y_max = pos[i, "y_max"]
        pushViewport(viewport(x = x_min, y = y_min, name = pos[i, "slice"],
            width = x_max - x_min, height = y_max - y_min,
            just = c("left", "bottom")))
        grid.rect()
        upViewport()
    }
    seekViewport("mat_a_heatmap_body_1_2")
    ht = ht_list@ht_list[["mat_a"]]
    m = ht@matrix

    i = 1
    j = 2
    row_order = ht@row_order_list[[i]]
    column_order = ht@column_order_list[[j]]
    nr = length(row_order)
    nc = length(column_order)
    grid.segments(1:nc/nc, rep(0, nc), 1:nc/nc, rep(1, nc), default.units = "npc",
        gp = gpar(col = "#888888", lty = 2))
    grid.segments(rep(0, nr), 1:nr/nr, rep(1, nr), 1:nr/nr, default.units = "npc",
        gp = gpar(col = "#888888", lty = 2))
    grid.rect(gp = gpar(fill = NA))

    grid.points(0.3, 0.8, pch = 16, size = unit(2, "mm"), gp = gpar(col = "blue"))
    ComplexHeatmap:::grid.text(gt_render("(a, b)", box_gp = gpar(fill = "white", col = NA)), 
        x = unit(0.3, "npc") + unit(2, "mm"), y = unit(0.8, "npc"),
        just = "left")

    grid.points(0, 0, pch = 16, size = unit(2, "mm"), gp = gpar(col = "red"))
    ComplexHeatmap:::grid.text(gt_render("(x<sub>1</sub>, y<sub>1</sub>)", 
        box_gp = gpar(fill = "white", col = NA)), 
        x = unit(0, "npc") + unit(2, "mm"), y = unit(0, "npc"),
        just = "left")
    grid.points(1, 1, pch = 16, size = unit(2, "mm"), gp = gpar(col = "red"))
    ComplexHeatmap:::grid.text(gt_render("(x<sub>2</sub>, y<sub>2</sub>)", 
        box_gp = gpar(fill = "white", col = NA)), 
        x = unit(1, "npc") + unit(2, "mm"), y = unit(1, "npc"),
        just = "left")

    ComplexHeatmap:::grid.text(gt_render("n<sub>r</sub> = 7", box_gp = gpar(fill = "white", col = NA)), 
        x = unit(1, "npc") + unit(1, "mm"), y = unit(0.5, "npc"),
        just = "left")

    ComplexHeatmap:::grid.text(gt_render("n<sub>c</sub> = 7", box_gp = gpar(fill = "white", col = NA)), 
        x = unit(0.5, "npc"), y = unit(1, "npc") + unit(1, "mm"),
        just = "bottom")
}, width = 6, height = 4)

print(plot_grid(p1, p2, nrow = 2, rel_heights = c(4, 4)))
```

#### Figure 8

```
library(ComplexHeatmap)
set.seed(111)
mat1 = matrix(rnorm(100), 10)
rownames(mat1) = colnames(mat1) = paste0("a", 1:10)
mat2 = matrix(sample(letters[1:10], 100, replace = TRUE), 10)
rownames(mat2) = colnames(mat2) = paste0("b", 1:10)

ht_list = Heatmap(mat1, name = "mat_a", row_km = 2, column_km = 2,
        top_annotation = HeatmapAnnotation(foo = anno_points(runif(10)))) +
    rowAnnotation(bar = anno_barplot(sample(10, 10))) +
    Heatmap(mat2, name = "mat_b")

ht_list = draw(ht_list) # not necessary, but recommended

df = selectPosition(ht_list)
```

#### Figure 9

```
library(ComplexHeatmap)
library(InteractiveComplexHeatmap)
library(shiny)

data(rand_mat) # simply a random matrix
ht1 = Heatmap(rand_mat, name = "mat",
    show_row_names = FALSE, show_column_names = FALSE)
ht1 = draw(ht1)

ui = fluidPage(
    h3("My first interactive ComplexHeatmap Shiny app"),
    p("This is an interactive heatmap visualization on a random matrix."),
    InteractiveComplexHeatmapOutput()
)
server = function(input, output, session) {
    makeInteractiveComplexHeatmap(input, output, session, ht1)
}
shinyApp(ui, server)
```

#### Figure 10

```
library(ComplexHeatmap)
library(InteractiveComplexHeatmap)
library(shiny)

data(rand_mat) # simply a random matrix
ht1 = Heatmap(rand_mat, name = "mat",
    show_row_names = FALSE, show_column_names = FALSE)
ht1 = draw(ht1)

set.seed(88)
mat2 = matrix(sample(letters[1:10], 100, replace = TRUE), 10)
ht2 = draw(Heatmap(mat2, name = "mat2"))

ui = fluidPage(
    h3("The first heatmap"),
    InteractiveComplexHeatmapOutput("heatmap_1", 
        height1 = 300, height2 = 300),
    hr(),
    h3("The second heatmap"),
    InteractiveComplexHeatmapOutput("heatmap_2", 
        height1 = 300, height2 = 300)
)
server = function(input, output, session) {
    makeInteractiveComplexHeatmap(input, output, session, ht1, "heatmap_1")
    makeInteractiveComplexHeatmap(input, output, session, ht2, "heatmap_2")
}
shinyApp(ui, server)
```

#### Figure 11

```
htShinyExample(10.1)
```

#### Figure 12

```
htShinyExample(5.5)
```

#### Figure 13

```
library(InteractiveComplexHeatmap)
library(shiny)
library(GetoptLong)
library(simplifyEnrichment)

mat = readRDS(system.file("extdata", "random_GO_BP_sim_mat.rds",
     package = "simplifyEnrichment"))
cl = binary_cut(mat)
ht = ht_clusters(mat, cl, draw_word_cloud = FALSE)

library(GO.db)
get_go_term = function(go_id) {
    term = suppressMessages(AnnotationDbi::select(GO.db, keys = go_id, columns = "TERM")$TERM)
    term[is.na(term)] = "NA"
    term
}

ui = fluidPage(
    InteractiveComplexHeatmapOutput(width1 = 700, height1 = 450, compact = TRUE,
        output_ui = htmlOutput("go_info"), title3 = NULL)
)

col_fun = ht@ht_list[[1]]@matrix_color_mapping@col_fun
library(GetoptLong)
click_action = function(df, output) {
    output[["go_info"]] = renderUI({
        if(!is.null(df)) {
            go_id1 = rownames(mat)[df$row_index]
            go_id2 = colnames(mat)[df$column_index]

            oe = try(term1 <- get_go_term(go_id1), silent = TRUE)
            if(inherits(oe, "try-error")) {
                term1 = ""
            }
            oe = try(term2 <- get_go_term(go_id2), silent = TRUE)
            if(inherits(oe, "try-error")) {
                term2 = ""
            }

            v = mat[go_id1, go_id2]
            col = col_fun(v)

            HTML(qq(
"<div style='padding:5px 10px;border:1px solid black; width:400px;'>
<h5>GO similarity</h5>
<p>@{sprintf('%.3f', v)}  <span style='background-color:@{col};width=10px;'>&nbsp;&nbsp;&nbsp;&nbsp;</span></p>
<h5>Row GO ID</h5>
<p><a href='http://amigo.geneontology.org/amigo/term/@{go_id1}' target='_blank'>@{go_id1}</a>: @{term1}</p>

<h5>Column GO ID</h5>
<p><a href='http://amigo.geneontology.org/amigo/term/@{go_id2}' target='_blank'>@{go_id2}</a>: @{term2}</p>
</div>"
))
        }
    })
}

server = function(input, output, session) {
    makeInteractiveComplexHeatmap(input, output, session, ht,
        click_action = click_action)
}

shinyApp(ui, server)
```

#### Figure 14

```
library(InteractiveComplexHeatmap)
library(ComplexHeatmap)
library(shiny)

ui = fluidPage(
    sliderInput("n_heatmap", label = "How many heatmaps?", 
        value = 1, min = 1, max = 5),
    InteractiveComplexHeatmapOutput()
)
generate_heatmap_list = function(n) {
    ht_list = NULL
    for(i in 1:n) {
        ht_list = ht_list + Heatmap(matrix(rnorm(100), 10), 
            name = paste0("mat_", i),
            column_title = paste0("heatmap_", i))
    }
    ht_list
}
server = function(input, output, session) {
    observe({
        ht_list = generate_heatmap_list(input$n_heatmap)
        makeInteractiveComplexHeatmap(input, output, session, ht_list)
    })
}
shiny::shinyApp(ui, server)
```

#### Figure 15

```
library(DESeq2)
library(airway)
data(airway)

dds = DESeqDataSet(airway, design = ~ dex)
dds = DESeq(dds)
interactivate(dds)
```

#### Session info

```
sessionInfo()
```

```
## R version 4.0.4 (2021-02-15)
## Platform: x86_64-apple-darwin17.0 (64-bit)
## Running under: macOS Big Sur 10.16
## 
## Matrix products: default
## BLAS:   /Library/Frameworks/R.framework/Versions/4.0/Resources/lib/libRblas.dylib
## LAPACK: /Library/Frameworks/R.framework/Versions/4.0/Resources/lib/libRlapack.dylib
## 
## locale:
## [1] C/en_US.UTF-8/C/C/C/C
## 
## attached base packages:
## [1] grid      stats     graphics  grDevices utils     datasets  methods  
## [8] base     
## 
## other attached packages:
##  [1] cowplot_1.1.1                         
##  [2] InteractiveComplexHeatmap_0.99.13.9000
##  [3] RColorBrewer_1.1-2                    
##  [4] dendextend_1.14.0                     
##  [5] GetoptLong_1.0.5                      
##  [6] circlize_0.4.12                       
##  [7] ComplexHeatmap_2.7.9.1002             
##  [8] knitr_1.31                            
##  [9] rmarkdown_2.7                         
## [10] colorout_1.2-2                        
## 
## loaded via a namespace (and not attached):
##  [1] httr_1.4.2          viridis_0.5.1       sass_0.3.1         
##  [4] jsonlite_1.7.2      viridisLite_0.3.0   foreach_1.5.1      
##  [7] bslib_0.2.4         shiny_1.6.0         assertthat_0.2.1   
## [10] highr_0.8           stats4_4.0.4        yaml_2.2.1         
## [13] pillar_1.5.1        glue_1.4.2          digest_0.6.27      
## [16] promises_1.2.0.1    gridtext_0.1.4      rvest_1.0.0        
## [19] colorspace_2.0-0    htmltools_0.5.1.1   httpuv_1.5.5       
## [22] clisymbols_1.2.0    pkgconfig_2.0.3     magick_2.7.1       
## [25] purrr_0.3.4         xtable_1.8-4        webshot_0.5.2      
## [28] scales_1.1.1        svglite_2.0.0       later_1.1.0.1      
## [31] tibble_3.1.0        generics_0.1.0      IRanges_2.24.1     
## [34] ggplot2_3.3.3       ellipsis_0.3.1      BiocGenerics_0.36.0
## [37] magrittr_2.0.1      crayon_1.4.1        mime_0.10          
## [40] evaluate_0.14       fansi_0.4.2         doParallel_1.0.16  
## [43] xml2_1.3.2          Cairo_1.5-12.2      tools_4.0.4        
## [46] GlobalOptions_0.1.2 lifecycle_1.0.0     matrixStats_0.58.0 
## [49] stringr_1.4.0       S4Vectors_0.28.1    munsell_0.5.0      
## [52] cluster_2.1.1       kableExtra_1.3.4    compiler_4.0.4     
## [55] jquerylib_0.1.3     systemfonts_1.0.1   rlang_0.4.10       
## [58] rstudioapi_0.13     iterators_1.0.13    rjson_0.2.20       
## [61] gtable_0.3.0        codetools_0.2-18    DBI_1.1.1          
## [64] markdown_1.1        R6_2.5.0            gridExtra_2.3      
## [67] dplyr_1.0.5         fastmap_1.1.0       utf8_1.2.1         
## [70] clue_0.3-58         shape_1.4.5         stringi_1.5.3      
## [73] parallel_4.0.4      Rcpp_1.0.6          vctrs_0.3.7        
## [76] png_0.1-7           tidyselect_1.1.0    xfun_0.22
```
