## Supplementary file 1 for "Make Interactive Complex Heatmaps in R": InteractiveComplexHeatmap_manuscript_code.md

## Figure 1

Figure A contains six panels:

### Figure 1A


```r
library(ComplexHeatmap)
```

```
## Loading required package: grid
```

```
## ========================================
## ComplexHeatmap version 2.7.9.1002
## Bioconductor page: http://bioconductor.org/packages/ComplexHeatmap/
## Github page: https://github.com/jokergoo/ComplexHeatmap
## Documentation: http://jokergoo.github.io/ComplexHeatmap-reference
## 
## If you use it in published research, please cite:
## Gu, Z. Complex heatmaps reveal patterns and correlations in multidimensional 
##   genomic data. Bioinformatics 2016.
## 
## The new InteractiveComplexHeatmap package can directly export static 
## complex heatmaps into an interactive Shiny app with zero effort. Have a try!
## 
## This message can be suppressed by:
##   suppressPackageStartupMessages(library(ComplexHeatmap))
## ========================================
```

```r
library(circlize)
```

```
## ========================================
## circlize version 0.4.12
## CRAN page: https://cran.r-project.org/package=circlize
## Github page: https://github.com/jokergoo/circlize
## Documentation: https://jokergoo.github.io/circlize_book/book/
## 
## If you use it in published research, please cite:
## Gu, Z. circlize implements and enhances circular visualization
##   in R. Bioinformatics 2014.
## 
## This message can be suppressed by:
##   suppressPackageStartupMessages(library(circlize))
## ========================================
```

```r
library(GetoptLong)

### figure A

set.seed(123)
nr1 = 4; nr2 = 8; nr3 = 6; nr = nr1 + nr2 + nr3
nc1 = 6; nc2 = 8; nc3 = 10; nc = nc1 + nc2 + nc3
mat = cbind(rbind(matrix(rnorm(nr1*nc1, mean = 1,   sd = 0.5), nr = nr1),
          matrix(rnorm(nr2*nc1, mean = 0,   sd = 0.5), nr = nr2),
          matrix(rnorm(nr3*nc1, mean = 0,   sd = 0.5), nr = nr3)),
    rbind(matrix(rnorm(nr1*nc2, mean = 0,   sd = 0.5), nr = nr1),
          matrix(rnorm(nr2*nc2, mean = 1,   sd = 0.5), nr = nr2),
          matrix(rnorm(nr3*nc2, mean = 0,   sd = 0.5), nr = nr3)),
    rbind(matrix(rnorm(nr1*nc3, mean = 0.5, sd = 0.5), nr = nr1),
          matrix(rnorm(nr2*nc3, mean = 0.5, sd = 0.5), nr = nr2),
          matrix(rnorm(nr3*nc3, mean = 1,   sd = 0.5), nr = nr3))
   )
mat = mat[sample(nr, nr), sample(nc, nc)] # random shuffle rows and columns
rownames(mat) = paste0("row", seq_len(nr))
colnames(mat) = paste0("column", seq_len(nc))

library(dendextend)
```

```
## 
## ---------------------
## Welcome to dendextend version 1.14.0
## Type citation('dendextend') for how to cite the package.
## 
## Type browseVignettes(package = 'dendextend') for the package vignette.
## The github page is: https://github.com/talgalili/dendextend/
## 
## Suggestions and bug-reports can be submitted at: https://github.com/talgalili/dendextend/issues
## Or contact: <>
## 
## 	To suppress this message use:  suppressPackageStartupMessages(library(dendextend))
## ---------------------
```

```
## 
## Attaching package: 'dendextend'
```

```
## The following object is masked from 'package:stats':
## 
##     cutree
```

```r
column_dend = as.dendrogram(hclust(dist(t(mat))))
column_dend = color_branches(column_dend, k = 3) # `color_branches()` returns a dendrogram object
column_dend = dendrapply(column_dend, function(d) {
    if(runif(1) > 0.5) attr(d, "nodePar") = list(cex = 0.8, pch = sample(20, 1), col = rand_color(1))
    return(d)
})
ht = Heatmap(mat, name = "mat", row_dend_width = unit(2, "cm"), cluster_columns = column_dend,
	column_names_rot = 45, row_split = rep(c("A", "B"), 9), row_km = 2, column_split = 3,
	top_annotation = HeatmapAnnotation(foo1 = 1:24, bar1 = anno_points(runif(24))),
    right_annotation = rowAnnotation(foo2 = 18:1, bar2 = anno_barplot(cbind(runif(18), runif(18)), gp = gpar(fill = 2:3), width = unit(2, "cm"))))

draw(ht, annotation_legend_list = list(Legend(title = "bar2", labels = c("group1", "group2"), legend_gp = gpar(fill = 2:3))),
	merge_legend = TRUE, padding = unit(c(5, 5, 5, 5), "mm"))
for(slice in 1:3) {
	decorate_annotation("bar1", slice = slice, {
		grid.lines(c(0, 1), c(0.5, 0.5), gp = gpar(lty = 2, col = "#AAAAAA"))
	})
}
```

![plot of chunk unnamed-chunk-1](figure/unnamed-chunk-1-1.png)


### figure 1B


```r
small_mat = mat[1:9, 1:9]
col_fun = colorRamp2(c(-2, 0, 2), c("green", "white", "red"))
ht = Heatmap(small_mat, name = "mat", col = col_fun,
    row_km = 2, column_km = 2,
    layer_fun = function(j, i, x, y, w, h, fill, slice_r, slice_c) {
        # restore_matrix() is explained after this chunk of code
        ind_mat = restore_matrix(j, i, x, y)
        for(ir in seq_len(nrow(ind_mat))) {
            # start from the second column
            for(ic in seq_len(ncol(ind_mat))[-1]) {
                ind1 = ind_mat[ir, ic-1] # previous column
                ind2 = ind_mat[ir, ic]   # current column
                v1 = small_mat[i[ind1], j[ind1]]
                v2 = small_mat[i[ind2], j[ind2]]
                if(v1 * v2 > 0) { # if they have the same sign
                    col = ifelse(v1 > 0, "violet", "blue")
                    grid.segments(x[ind1], y[ind1], x[ind2], y[ind2],
                        gp = gpar(col = col, lwd = 2))
                    grid.points(x[c(ind1, ind2)], y[c(ind1, ind2)], 
                        pch = 16, gp = gpar(col = col), size = unit(4, "mm"))
                }
            }
        }
        if(slice_r != slice_c) {
            grid.rect(gp = gpar(lwd = 2, fill = "transparent"))
        }
    }
)
draw(ht, padding = unit(c(5, 5, 5, 5), "mm"))
```

![plot of chunk unnamed-chunk-2](figure/unnamed-chunk-2-1.png)


### Figure C


```r
image_png = sample(dir("~/project/development/ComplexHeatmap-reference/IcoMoon-Free-master/PNG/64px", full.names = TRUE), 20)


set.seed(123)
n = 7
m = matrix(rnorm(100*n), nc = 100)
m2 = matrix(rnorm(100*n), nc = 100)

ha1 = rowAnnotation(
	"anno_simple" = runif(n),
	"anno_image" = anno_image(image_png[1:n], gp = gpar(col = "black"), space= unit(2, "mm"), width = unit(1, "cm")),
	"anno_points" = anno_points(matrix(runif(2*n), nc = 2), pch = 1:2, gp = gpar(col = 2:3), width = unit(2, "cm")),
	"anno_lines" = anno_lines(cbind(runif(n), runif(n)), gp = gpar(col = 2:3, lty = 1:2), width = unit(2, "cm")),
	"anno_barplot" = anno_barplot(cbind(runif(n), runif(n)), gp = gpar(fill = 2:3, col = 2:3), width = unit(2, "cm")),
	"anno_boxplot" = anno_boxplot(m, gp = gpar(fill = 1:n), width = unit(2, "cm")),
	"anno_text" = anno_text(gt_render(qq("row<sub>@{1:n}</sub>", collapse = FALSE), align_widths = TRUE, halign = 0.5, 
                        r = unit(4, "pt"),
                        padding = unit(c(2, 8, 2, 8), "pt")), 
                    gp = gpar(box_col = "blue", box_lwd = 2, col = 1:10, fontfamily = "Times"), 
                    just = "right", 
                    location = unit(1, "npc")
    ),
    "anno_histogram" = anno_histogram(m2, gp = gpar(fill = 1:n), border = TRUE, width = unit(3.5, "cm")),
	"anon_violin" = anno_density(m, type = "violin", gp = gpar(fill = 1:10, col = c("white", rep("black", 7), "white", "black")), width = unit(3.5, "cm")),
	show_legend = FALSE, simple_anno_size = unit(1, "cm"), gap = unit(2, "mm"), show_annotation_name = FALSE
)
```

```
## Loading required namespace: gridtext
```

```r
m = matrix(rnorm(1500), nc = 15)
lt1 = apply(m, 2, function(x) data.frame(density(x)[c("x", "y")]))

lt2 = lapply(1:15, function(x) cumprod(1 + runif(1000, -x/100, x/100)) - 1)

# 20 rows
ha3 = rowAnnotation(
	"anno_joyplot" = anno_joyplot(lt1, width = unit(4, "cm"), gp = gpar(fill = 1:10), transparency = 0.75, scale = 2),
	"anno_horizon" = anno_horizon(lt2),
    gap = unit(2, "mm"), show_annotation_name = FALSE
)

grid.newpage()
grid.rect(gp = gpar(fill = "white", col = NA))
pushViewport(viewport(width = ComplexHeatmap:::width(ha1) + ComplexHeatmap:::width(ha3)))

pushViewport(viewport(x = 0, y = unit(1, "npc") - unit(5, "mm"), width = ComplexHeatmap:::width(ha1), height = unit(1, "npc") - unit(2, "cm"), just = c("left", "top")))
draw(ha1)
popViewport()

pushViewport(viewport(x = ComplexHeatmap:::width(ha1) + unit(2, "mm"), y = unit(1, "npc") - unit(5, "mm"), width = ComplexHeatmap:::width(ha3) - unit(2, "mm"), height = unit(1, "npc") - unit(2, "cm"), just = c("left", "top")))
draw(ha3)
popViewport()

popViewport()
```

![plot of chunk unnamed-chunk-3](figure/unnamed-chunk-3-1.png)


### figure D


```r
res_list = readRDS(url("http://jokergoo.github.io/supplementary/ComplexHeatmap-supplementary1-4/supplS3_methylation/meth.rds"))
type = res_list$type
mat_meth = res_list$mat_meth
mat_expr = res_list$mat_expr
direction = res_list$direction
cor_pvalue = res_list$cor_pvalue
gene_type = res_list$gene_type
anno_gene = res_list$anno_gene
dist = res_list$dist
anno_enhancer = res_list$anno_enhancer

column_tree = hclust(dist(t(mat_meth)))
column_order = column_tree$order

library(RColorBrewer)
meth_col_fun = colorRamp2(c(0, 0.5, 1), c("blue", "white", "red"))
direction_col = c("hyper" = "red", "hypo" = "blue")
expr_col_fun = colorRamp2(c(-2, 0, 2), c("green", "white", "red"))
pvalue_col_fun = colorRamp2(c(0, 2, 4), c("white", "white", "red"))
gene_type_col = structure(brewer.pal(length(unique(gene_type)), "Set3"), 
    names = unique(gene_type))
anno_gene_col = structure(brewer.pal(length(unique(anno_gene)), "Set1"), 
    names = unique(anno_gene))
dist_col_fun = colorRamp2(c(0, 10000), c("black", "white"))
enhancer_col_fun = colorRamp2(c(0, 1), c("white", "orange"))

ht_opt(
    legend_title_gp = gpar(fontsize = 8, fontface = "bold"), 
    legend_labels_gp = gpar(fontsize = 8), 
    heatmap_column_names_gp = gpar(fontsize = 8),
    heatmap_column_title_gp = gpar(fontsize = 10),
    heatmap_row_title_gp = gpar(fontsize = 8)
)

ha = HeatmapAnnotation(type = type, 
    col = list(type = c("Tumor" = "pink", "Control" = "royalblue")),
    annotation_name_side = "left")
ha2 = HeatmapAnnotation(type = type, 
    col = list(type = c("Tumor" = "pink", "Control" = "royalblue")), 
    show_legend = FALSE)

ht_list = Heatmap(mat_meth, name = "methylation", col = meth_col_fun,
    column_order= column_order,
    top_annotation = ha, column_title = "Methylation") +
    Heatmap(direction, name = "direction", col = direction_col) +
    Heatmap(mat_expr[, column_tree$order], name = "expression", 
        col = expr_col_fun, 
        column_order = column_order, 
        top_annotation = ha2, column_title = "Expression") +
    Heatmap(cor_pvalue, name = "cor_p", col = pvalue_col_fun) +
    Heatmap(gene_type, name = "gene type", col = gene_type_col) +
    Heatmap(anno_gene, name = "anno_gene", col = anno_gene_col) +
    Heatmap(dist, name = "dist_tss", col = dist_col_fun) +
    Heatmap(anno_enhancer, name = "anno_enhancer", col = enhancer_col_fun, 
        cluster_columns = FALSE, column_title = "Enhancer")
```

```
## Warning: The input is a data frame, convert it to the matrix.
```

```r
draw(ht_list, row_km = 2, row_split = direction,
    column_title = "Comprehensive correspondence between methylation, expression and other genomic features", 
    column_title_gp = gpar(fontsize = 12, fontface = "bold"), 
    merge_legends = TRUE, padding = unit(c(5, 15, 5, 15), "mm"))
```

![plot of chunk unnamed-chunk-4](figure/unnamed-chunk-4-1.png)

```r
ht_opt(RESET = TRUE)
```


## Figure 2


```r
library(ComplexHeatmap)
set.seed(111)
mat1 = matrix(rnorm(100), 10)
rownames(mat1) = colnames(mat1) = paste0("a", 1:10)
mat2 = matrix(sample(letters[1:10], 100, replace = TRUE), 10)
rownames(mat2) = colnames(mat2) = paste0("b", 1:10)

ht_list = Heatmap(mat1, name = "mat_a", row_km = 2, column_km = 2,
        top_annotation = HeatmapAnnotation(foo = anno_points(runif(10)))) +
    rowAnnotation(bar = anno_barplot(sample(10, 10))) +
    Heatmap(mat2, name = "mat_b")

ht_list = draw(ht_list) # not necessary, but recommended

library(InteractiveComplexHeatmap)
htShiny(ht_list)
```


## Figure 3


```r
library(InteractiveComplexHeatmap)
library(cola)  # from Bioconductor
data(golub_cola)
get_signatures(golub_cola["ATC:skmeans"], k = 3) # this makes the heatmap
ht_shiny()
```


## Figure 4


```r
htShinyExample(4.2)
```

## Figure 5

### figure 5A


```r
htShinyExample(2.5)
```

### figure 5B


```r
htShinyExample(2.6)
```

### figure 5C


```r
htShinyExample(2.4)
```

## Figure 6

### figure 6A


```r
htShinyExample(2.2)
```

### figure 5B


```r
htShinyExample(2.3)
```

### figure 5C


```r
htShinyExample(2.1)
```

### figure 5D


```r
htShinyExample(3.1)
```

## Figure 8


```r
library(ComplexHeatmap)
set.seed(111)
mat1 = matrix(rnorm(100), 10)
rownames(mat1) = colnames(mat1) = paste0("a", 1:10)
mat2 = matrix(sample(letters[1:10], 100, replace = TRUE), 10)
rownames(mat2) = colnames(mat2) = paste0("b", 1:10)

ht_list = Heatmap(mat1, name = "mat_a", row_km = 2, column_km = 2,
        top_annotation = HeatmapAnnotation(foo = anno_points(runif(10)))) +
    rowAnnotation(bar = anno_barplot(sample(10, 10))) +
    Heatmap(mat2, name = "mat_b")

ht_list = draw(ht_list) # not necessary, but recommended

df = selectPosition(ht_list)
```

## Figure 9


```r
library(ComplexHeatmap)
library(InteractiveComplexHeatmap)
library(shiny)

data(rand_mat) # simply a random matrix
ht1 = Heatmap(rand_mat, name = "mat",
    show_row_names = FALSE, show_column_names = FALSE)
ht1 = draw(ht1)

ui = fluidPage(
    h3("My first interactive ComplexHeatmap Shiny app"),
    p("This is an interactive heatmap visualization on a random matrix."),
    InteractiveComplexHeatmapOutput()
)
server = function(input, output, session) {
    makeInteractiveComplexHeatmap(input, output, session, ht1)
}
shinyApp(ui, server)
```


## Figure 10


```r
library(ComplexHeatmap)
library(InteractiveComplexHeatmap)
library(shiny)

data(rand_mat) # simply a random matrix
ht1 = Heatmap(rand_mat, name = "mat",
    show_row_names = FALSE, show_column_names = FALSE)
ht1 = draw(ht1)

set.seed(88)
mat2 = matrix(sample(letters[1:10], 100, replace = TRUE), 10)
ht2 = draw(Heatmap(mat2, name = "mat2"))

ui = fluidPage(
    h3("The first heatmap"),
    InteractiveComplexHeatmapOutput("heatmap_1", 
        height1 = 300, height2 = 300),
    hr(),
    h3("The second heatmap"),
    InteractiveComplexHeatmapOutput("heatmap_2", 
        height1 = 300, height2 = 300)
)
server = function(input, output, session) {
    makeInteractiveComplexHeatmap(input, output, session, ht1, "heatmap_1")
    makeInteractiveComplexHeatmap(input, output, session, ht2, "heatmap_2")
}
shinyApp(ui, server)
```


## Figure 11


```r
htShinyExample(10.1)
```


## Figure 12


```r
htShinyExample(5.5)
```

## Figure 13


```r
library(GetoptLong)
suppressPackageStartupMessages(library(simplifyEnrichment))

mat = readRDS(system.file("extdata", "random_GO_BP_sim_mat.rds",
     package = "simplifyEnrichment"))
cl = binary_cut(mat)
ht = ht_clusters(mat, cl, draw_word_cloud = FALSE)

suppressPackageStartupMessages(library(GO.db))
get_go_term = function(go_id) {
    term = suppressMessages(AnnotationDbi::select(GO.db, keys = go_id, columns = "TERM")$TERM)
    term[is.na(term)] = "NA"
    term
}

ui = fluidPage(
    InteractiveComplexHeatmapOutput(width1 = 700, height1 = 450, compact = TRUE,
        output_ui = htmlOutput("go_info"), title3 = NULL)
)

col_fun = ht@ht_list[[1]]@matrix_color_mapping@col_fun
library(GetoptLong)
click_action = function(df, output) {
    output[["go_info"]] = renderUI({
        if(!is.null(df)) {
            go_id1 = rownames(mat)[df$row_index]
            go_id2 = colnames(mat)[df$column_index]

            oe = try(term1 <- get_go_term(go_id1), silent = TRUE)
            if(inherits(oe, "try-error")) {
                term1 = ""
            }
            oe = try(term2 <- get_go_term(go_id2), silent = TRUE)
            if(inherits(oe, "try-error")) {
                term2 = ""
            }

            v = mat[go_id1, go_id2]
            col = col_fun(v)

            HTML(qq(
"<div style='padding:5px 10px;border:1px solid black; width:400px;'>
<h5>GO similarity</h5>
@{sprintf('%.3f', v)}  <span style='background-color:@{col};width=10px;'>&nbsp;&nbsp;&nbsp;&nbsp;</span>
<h5>Row GO ID</h5>
<p><a href='http://amigo.geneontology.org/amigo/term/@{go_id1}' target='_blank'>@{go_id1}</a>: @{term1}</p>

<h5>Column GO ID</h5>
<p><a href='http://amigo.geneontology.org/amigo/term/@{go_id2}' target='_blank'>@{go_id2}</a>: @{term2}</p>
</div>"
))
        }
    })
}

server = function(input, output, session) {
    makeInteractiveComplexHeatmap(input, output, session, ht,
        click_action = click_action)
}

shinyApp(ui, server)
```


## Figure 14


```r
ui = fluidPage(
    sliderInput("n_heatmap", label = "How many heatmaps?", 
        value = 1, min = 1, max = 5),
    InteractiveComplexHeatmapOutput()
)
generate_heatmap_list = function(n) {
    ht_list = NULL
    for(i in 1:n) {
        ht_list = ht_list + Heatmap(matrix(rnorm(100), 10), 
            name = paste0("mat_", i),
            column_title = paste0("heatmap_", i))
    }
    ht_list
}
server = function(input, output, session) {
    observe({
        ht_list = generate_heatmap_list(input$n_heatmap)
        makeInteractiveComplexHeatmap(input, output, session, ht_list)
    })
}
shiny::shinyApp(ui, server)
```


## Figure 15


```r
library(DESeq2)
library(airway)
data(airway)

dds = DESeqDataSet(airway, design = ~ dex)
dds = DESeq(dds)
interactivate(dds)
```
